# Selective Vulnerability of Parvocellular Oxytocin Neurons in Social Dysfunction

**DOI:** 10.1101/2023.12.02.569733

**Authors:** Masafumi Tsurutani, Teppei Goto, Mitsue Hagihara, Satsuki Irie, Kazunari Miyamichi

**Affiliations:** Laboratory for Comparative Connectomics, RIKEN Center for Biosystems Dynamics Research, Kobe, Hyogo 650-0047, Japan; Graduate School of Biostudies, Kyoto University, Kyoto, Kyoto 606-8501, Japan

## Abstract

Selective vulnerability offers a conceptual framework for understanding neurodegenerative disorders, such as Parkinson’s disease, where specific neuronal types are selectively affected while adjacent ones are spared. The applicability of this framework to neurodevelopmental disorders remains uncertain, particularly those characterized by atypical social behaviors such as autism spectrum disorder. Here, employing a single-cell transcriptome analysis in mice, we show that an embryonic disturbance known to induce social dysfunction preferentially impairs gene expressions crucial for neural functions in parvocellular oxytocin (OT) neurons—a subtype linked to social rewards—while neighboring cell types experience a lesser impact. Chemogenetic stimulation of OT neurons at the neonatal stage ameliorated social deficits in early adulthood, concurrent with a cell-type-specific sustained recovery of the pivotal gene expressions within parvocellular OT neurons. Collectively, our data shed light on the transcriptomic selective vulnerability within the hypothalamic social behavioral center and provide a potential therapeutic target through specific neonatal neurostimulation.

## Introduction

The applicability of the selective vulnerability framework^1^ to neurodevelopmental disorders (NDDs) remains uncertain. The pathogenesis of NDDs is closely associated with fetal genetic factors and maternal/environmental influences, such as maternal immune activation, gut microbiota, and medications administered during fetal brain development^2–5^. As these factors exert systemic effects on the entire nervous system, the precise mechanisms underlying the pathogenesis of NDDs remain largely elusive. For example, despite rapid advancements in our understanding of neural circuits that regulate social behaviors in rodents^6, 7^, it remains unclear whether specific neural cell types are selectively affected in pathological conditions that model NDDs. To fill this knowledge gap, we aim to focus on the OT neurons in the paraventricular hypothalamus (PVH^OT^ neurons) in mouse models that exhibit social dysfunction.

Decades of rodent studies have highlighted the significance of the OT system in the typical development of social behaviors, bond formation, and parental behaviors^6, 8, 9^. Numerous brain regions express OT receptors^10^, emphasizing its widespread functions throughout the brain. Among the population of OT neurons, magnocellular PVH^OT^ neurons exhibit bifurcated axonal projections to the posterior pituitary and various forebrain structures^11^. In contrast, parvocellular PVH^OT^ neurons selectively project to the central brain, including the ventral tegmental area and substantia nigra, thereby modulating the dopamine system and contributing to social reward^12–14^. Additionally, parvocellular PVH^OT^ neurons innervate the hindbrain and spinal cord, exerting diverse neuromodulatory effects on emotions, appetite, and pain^11, 15^.

Impairment of the OT system has been extensively documented in various genetic and environmental rodent models of social dysfunction. For instance, reductions in OT immunoreactivity or *OT* mRNA expression within the PVH have been observed in genetic mutants of the *Shank3b* gene^5, 16^, *Maged1* gene^17^, *Necdin* gene^18^, the progeny of mothers consuming a high-fat diet^4^, and rats prenatally exposed to valproic acid (VPA)^19, 20^. However, these studies have not discerned whether the observed reduction stems from a decline in OT expression or a loss of PVH^OT^ neurons at the cellular level. Furthermore, the potential impacts on distinct cell types of PVH^OT^ neurons remain unknown. A recent seminal study employing single-cell RNA sequencing (RNA-seq) has unveiled distinct transcriptomic signatures of magnocellular and parvocellular PVH^OT^ neurons, with the latter displaying an enriched expression of ASD risk factor genes^21^ and playing a more significant role in social reward^12^. This lets us hypothesize that parvocellular PVH^OT^ neurons may be more susceptible to disruptions caused by embryonic factors that induce social dysfunctions.

## Result

### Histochemical analyses of OT ligands in parvocellular PVH^OT^ neurons

We first examined a mouse model of prenatal exposure to VPA^19, 22^. We confirmed that VPA-treated male *OT-Cre* mice^23^ on the C57BL/6 background exhibited decreased sociability, as assessed by the three-chamber test (Fig. 1a). In contrast to control mice with prenatal saline exposure, which spent significantly more time investigating a cage containing an unfamiliar mouse compared to a cage with a non-animal object, the VPA-treated group failed to show this preference (Fig. 1b). We then performed intravenous Fluoro-Gold injection to selectively visualize magnocellular but not parvocellular PVH^OT^ neurons based on the fact that only magnocellular OT neurons send axonal projections outside the blood-brain barrier^12^ (Fig. 1c). Additionally, we utilized *OT-Cre*; *Ai9* (tdTomato Cre reporter mouse line^24^) double knockin mice to label OT neurons genetically regardless of OT ligand expression at the time of analysis. We found that the total number of tdTomato-labeled cells was unaltered in VPA-treated mice (Fig. 1d). Consistent with previous research^12^, magnocellular PVH^OT^ neurons were located in the anterior part of the PVH, whereas parvocellular PVH^OT^ neurons were located in the posterior part (Fig. 1e, f). The number of OT-expressing cells, as detected by anti-OT antibodies, was selectively reduced in the parvocellular PVH^OT^ neurons (Fig. 1g, top). Quantitative analysis of fluorescent intensity indicated a significant reduction of OT ligand expression in both types of PVH^OT^ neurons, with a more pronounced reduction in the parvocellular PVH^OT^ neurons (Fig. 1g, bottom). These findings exclude the possibility of a cellular loss of PVH^OT^ neurons in VPA-treated mice and demonstrate a reduction in OT expression at the protein level, preferentially affecting parvocellular PVH^OT^ neurons.

**Fig. 1:**
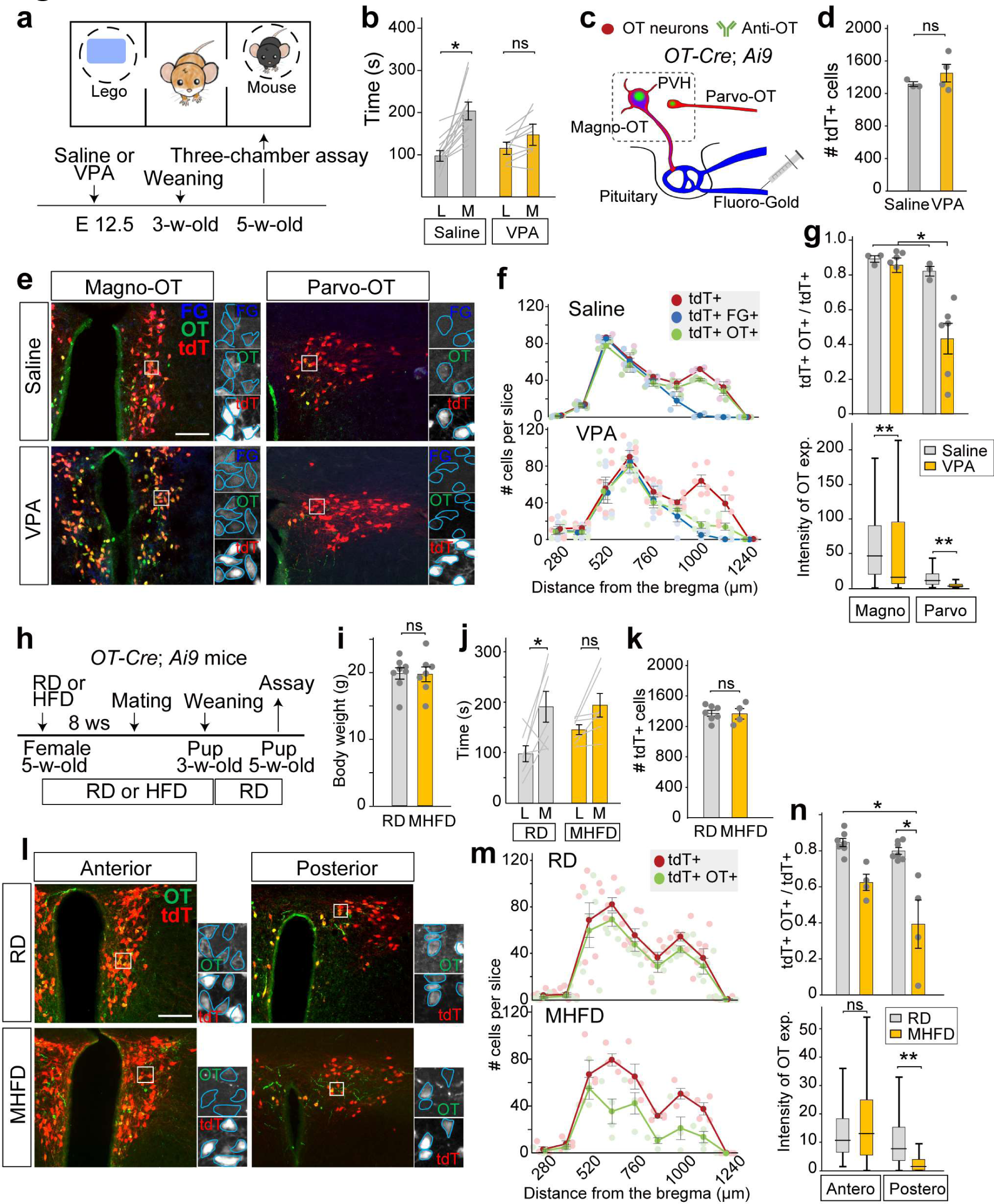
Reduction of OT ligand expression in parvocellular PVHOT neurons. **a**–**g**, Behavior and OT ligand expression of VPA-treated mice. **a**, Timeline of the experiment (bottom) and schematic diagram of the three-chamber assay (top). E, embryonic day. **b**, Duration of mice exploring a non-social object (Lego, L) and an unfamiliar male mouse (M). N = 14 for the saline and N = 9 for the VPA group. *, p < 0.05 by the Wilcoxon rank-sum test. **c**, Schematic diagrams depicting the labeling strategy. **d**, Number of tdT+ cells in the PVH. No difference was detected by the Wilcoxon rank-sum test. **e**, Typical coronal sections of the PVH showing tdTomato (tdT+), anti-OT staining (OT+), and Fluoro-Gold (FG+). Scale bar, 200 μm. **f**, Number of tdT+, tdT+ FG+, and tdT+ OT+ cells per slice. N = 3 and 6 for the saline and VPA groups. **g**, (Top) Fraction of tdT+ OT+ cells over tdT+ cells. *, p < 0.05 by two-way ANOVA with the post-hoc Tukey–Kramer test. (Bottom) Fluorescent intensity of anti-OT staining. **, p < 0.01 by the Wilcoxon rank-sum test with Bonferroni correction. **h–n**, Behavior and OT ligand expression of HMFD group. **h**, Timeline of the experiment. RD, regular diet; HFD, high-fat diet. **i**, Body weight at 5 weeks of age. No difference was detected by the Wilcoxon rank-sum test. **j**, Duration of mice investigating a Lego block (L) and an unfamiliar male mouse (M). N = 7 each. *, p < 0.05 by the Wilcoxon rank-sum test. **k**, Number of tdT+ cells. No difference was detected by the Wilcoxon rank-sum test. **l**, Typical coronal sections of the PVH showing tdTomato (tdT+) and anti-OT staining (OT+). Scale bar, 200 μm. **m**, Number of tdT+ and tdT+ OT+ cells per slice. N = 7 and 4 for the RD and MHFD groups. **n**, (Top) Fraction of tdT+ OT+ cells over tdT+ cells. *, p < 0.05 by two-way ANOVA with the post-hoc Tukey–Kramer test. (Bottom) Fluorescent intensity of anti-OT staining. **, p < 0.01 by the Wilcoxon rank-sum test with Bonferroni correction. Error bars, SEM.

To examine the generality of these findings, we conducted similar experiments in mice born to mothers that had been fed a high-fat diet, which we refer to as the maternal high-fat diet (MHFD) group for simplicity (Fig. 1h). Consistent with a previous study^4^, the body weight of the MHFD group did not differ from that of control mice born to mothers that had been fed a regular diet (RD, Fig. 1i), and the MHFD group showed reduced sociability (Fig. 1j). Although Fluoro-Gold labeling of magnocellular PVH^OT^ neurons was not effective in the MHFD group for unknown reasons, we were able to show a significant reduction of anti-OT immunostaining along the entire anterior–posterior axis of the PVH, without affecting the total number of *Ai9*-labeled cells (Fig. 1k–m). Focusing on the posterior part of the PVH where the parvocellular PVH^OT^ neurons exist, we observed a significant decrease in the number of OT ligand-expressing cells and fluorescent intensity in individual cells in the MHFD compared with the RD control group (Fig. 1n). These results indicate that two independent mouse models exhibiting atypical sociability due to exogenous or maternal factors commonly display reduced OT ligand expression in parvocellular PVH^OT^ neurons.

### Aberrant gene expression in parvocellular PVH^OT^ neurons

Next, we aimed to investigate whether the reduction in OT expression occurred at the mRNA level and, if so, whether aberrant gene expression was specific to the *OT* gene or more widespread. Additionally, we aimed to assess the impact on other neural cell types located within or near the PVH. To address these inquiries, we surgically dissected a hypothalamic region containing the PVH from VPA-treated *OT-Cre*; *Ai9* male mice, as well as from the control group prenatally exposed to saline. Single nucleus (sn) RNA-seq profiles were obtained using the 10X Genomics Chromium platform for 10,060 cells in the VPA-treated group and 3750 cells in the saline-treated groups that met the quality control standards (Methods). We combined the sequencing data from both groups and employed unsupervised graph-based clustering using Cell Ranger^25^ and Seurat^26^ to classify 30 clusters of excitatory neurons expressing *vesicular glutamate transporter type 2* (*vGluT2*) (Fig. 2a, b), following a more general categorization (Extended Data Figs. 1 and 2). The VPA- and saline-treated groups were intermingled within these excitatory clusters, indicating that VPA did not affect the overall transcriptomic signature (Fig. 2a and Extended Data Fig. 1d).

**Fig. 2:**
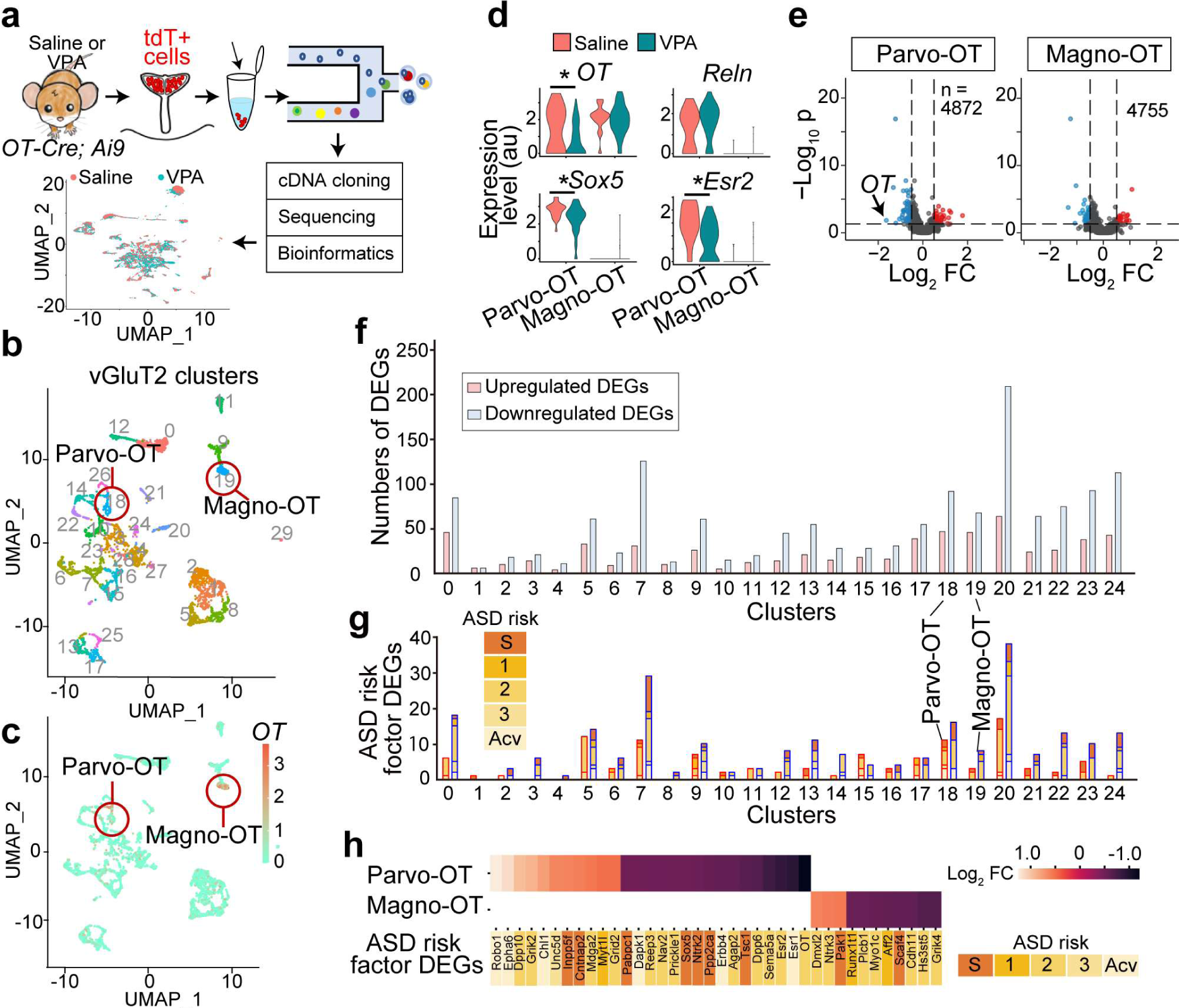
Transcriptomic profiles of PVH^OT^ neurons in VPA-treated mice. **a**, Schematics of single nucleus RNA-seq data collection and analysis. tdT, tdTomato from the *Ai9* allele. Universal manifold approximation and projection (UMAP) representation of all nucleus data are also shown. **b**, **c**, UMAP representation of *vGluT2*+ excitatory clusters (**b**) and *OT* gene expression, with a color scale showing log-normalized expression (**c**). **d**, Violin plots of *OT*, and three genetic markers (*Sox5*, *Reln*, and *Esr2* genes) to distinguish parvocellular (cluster 18) and magnocellular (cluster 19) PVH^OT^ neurons. *, p < 0.05 by the Wilcoxon rank-sum test. **e**, Volcano plots representing DEGs (upregulated, red dots; downregulated, blue dots) in the parvocellular and magnocellular PVH^OT^ neurons. The X-axis represents the log2-transformed fold-change ratios, and the Y-axis represents the log10-transformed p-value. The number (n) of genes analyzed is indicated in the panel. **f**, The numbers of upregulated (red) and downregulated (blue) DEGs within the *vGluT2+* clusters. The clusters 25–29 were excluded from this analysis as the number of nuclei was too small (less than 57). **g**, The number of ASD risk factor DEGs within the upregulated (red frame) and downregulated (blue frame) DEGs. **h**, Heatmaps of the log2-transformed fold change for ASD risk factor DEGs found within parvocellular and magnocellular PVH^OT^ neurons. The color codes for graphs (**g**) and gene names (**h**) show the rank of ASD risk^21^. For more data, see Extended Data Figs. 1–3 and Supplementary Table 1.

OT-positive clusters were readily identifiable owing to their distinctive expression of *OT* genes (Fig. 2b, c). Based on an analysis of gene expression, including *Sox5*, *estrogen receptor type 2* (*Esr2*), and *Reelin* genes, as outlined in a previous study^12^ and our own *in situ* hybridization (ISH) data (Extended Data Fig. 3), we were able to identify the parvocellular (cluster 18) and magnocellular (cluster 19) PVH^OT^ neurons (Fig. 2c, d). Differentially expressed genes (DEGs) between the VPA-treated and control groups were defined as those exhibiting a >1.4-fold change and satisfying a false discovery rate criterion of < 0.05 (Fig. 2e, Extended Data Fig. 2b). Notably, the expression of the *OT* gene itself was significantly reduced in the parvocellular PVH^OT^ neurons of VPA-treated compared with the saline-treated control mice (Fig. 2d, e), suggesting that the observed downregulation of OT ligand expression (Fig. 1e–g) is a result of decreased mRNA expression. We found varying numbers of both upregulated and downregulated DEGs among the top 25 *vGluT2*-positive (+) clusters (Fig. 2f). Additionally, we identified variable numbers of DEGs that were previously designated as high-confidence ASD risk factor genes^21^ within each *vGluT2*+ cluster (Fig. 2g). Importantly, each cluster displayed a distinct set of ASD risk factor DEGs, with only a small fraction of genes common in two or more clusters (Supplementary Table 1). For instance, the magnocellular and parvocellular PVH^OT^ neurons showed completely non-overlapping sets of ASD risk factor DEGs (Fig. 2h). These data demonstrate that VPA affects not only the *OT* gene in parvocellular PVH^OT^ neurons but also numerous other genes, including diverse ASD risk factor genes, in broad cell clusters near and within the PVH.

We then conducted Gene Ontology (GO) analysis^27, 28^ for the DEGs found within the top 25 *vGluT2*+ clusters (Fig. 3a). Among them, the parvocellular PVH^OT^ neurons (cluster 18) exhibited the highest number of significantly associated GO terms for both upregulated and downregulated DEGs (Fig. 3a, Extended Data Fig. 4, and Supplementary Table 2). Specifically, we observed an enrichment of DEGs involved in synaptic functions, behavioral regulations, and intracellular signal transduction in the parvocellular PVH^OT^ neurons, whereas the DEGs in the arginine vasopressin (AVP; cluster 11) or magnocellular PVH^OT^ (cluster 19) neurons displayed little or no association with these functions. Moreover, when subjecting the downregulated DEGs to pathway analysis^29^, we found that parvocellular PVH^OT^ neurons were significantly affected in the largest number of signaling pathways with diverse biological functions (Fig. 3b, Extended Data Fig. 3). These pathways involved the phosphatidylinositol 3-kinase (PI3K)/protein kinase B (Akt) signaling pathway (Fig. 3c), which is relevant to the subcellular integration of the synaptic neurotransmission and neural plasticity in the brain^30^. ASD risk factor genes were predominantly enriched in this pathway (Extended Data Fig. 4b). In contrast, pathways significantly associated with the upregulated DEGs were more enriched in other cell clusters (Extended Data Fig 4c). To validate the DEGs using an independent method, we visualized the expression of some of the downregulated ASD risk factor DEGs found in the parvocellular PVH^OT^ neurons *in vivo* through ISH. All eight genes that we examined exhibited a significant reduction in mRNA expression (Extended Data Fig. 5). Notably, the downregulation of PI3K/Akt pathway-related genes, such as *Erbb4*, *Ntrk2*, and *Tsc1*, was confirmed. These findings suggest that parvocellular PVH^OT^ neurons are not distinct based on the numbers of DEGs or ASD risk factor DEGs, but rather they are distinguished by a unique impact on crucial neural functions and pathways in VPA-treated mice.

**Fig. 3:**
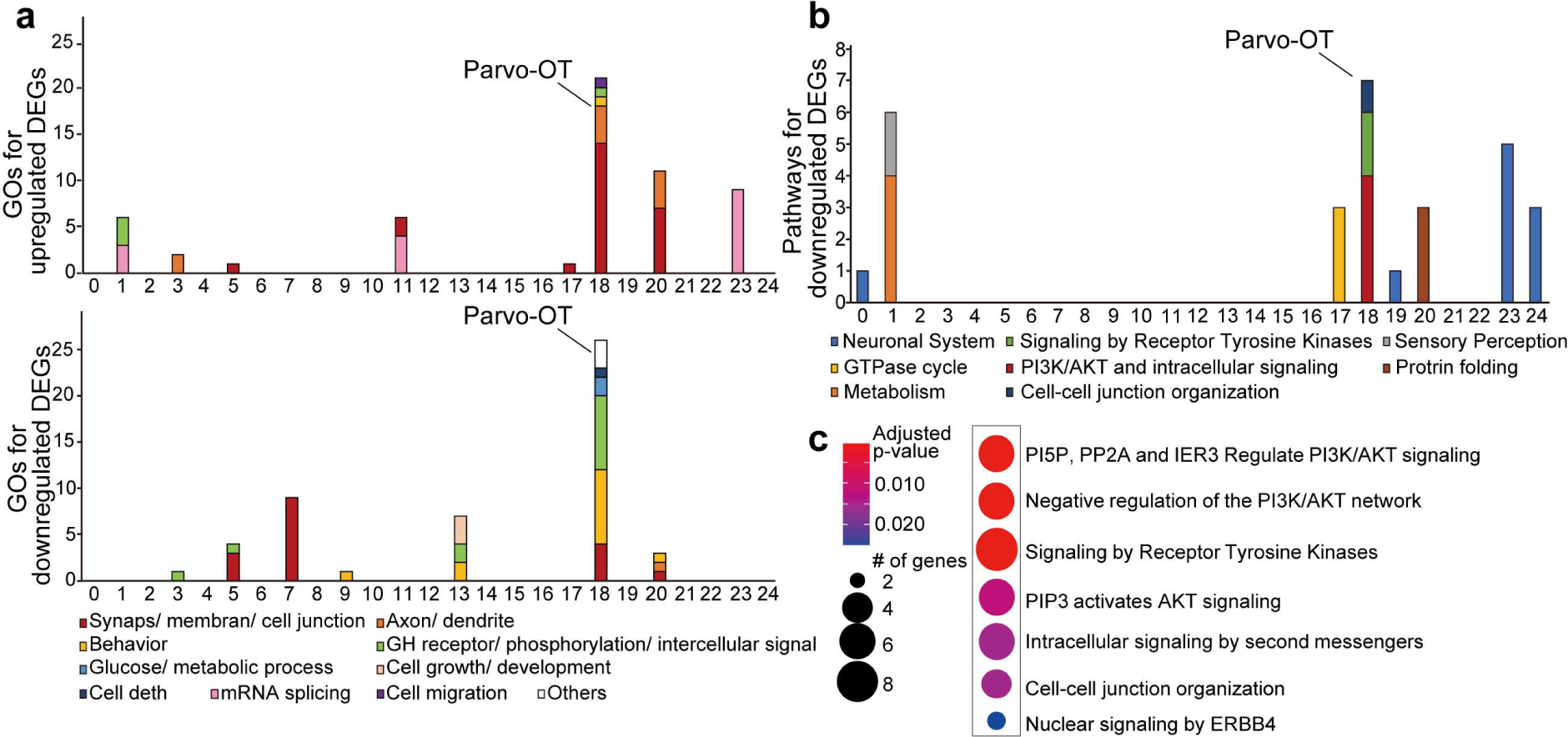
Gene Ontology (GO) and pathway analyses of DEGs. **a**, **b**, The numbers of GO terms (**a**) and pathways (**b**) that show significant association (GOs, p < 0.01 and pathways, p < 0.05) with DEGs within the top 25 *vGluT2+* clusters. The full list of GO terms and pathways for each cluster is provided in Supplementary Tables 2 and 3. Pathway analysis for the upregulated DEGs is shown in Extended Data Fig. 4b. **c**, Heatmap of p-values for the pathway analysis using downregulated DEGs in the parvocellular PVH^OT^ neurons. For more data, see Extended Data Figs. 4–6. For abbreviations and details of GO terms and pathways, see Supplementary Tables 2 and 3.

In addition to the excitatory clusters, our data also suggested a potential abnormality in gene expression profiles in the inhibitory neurons within or near the PVH. Unsupervised graph-based clustering revealed 17 clusters positive for the *glutamate decarboxylase type 2* (*GAD2*) gene (Extended Data Fig. 6a, b). We found varying numbers of both upregulated and downregulated DEGs, and ASD risk-factor DEGs, in the *GAD2*-positive clusters (Extended Data Fig 6c, d). Among them, cluster 10 (positive for *estrogen receptor type 1* gene, or *Esr1*) and cluster 11 (positive for *thyrotropin-releasing hormone receptor* gene, or *Trhr*) showed particular enrichments of diverse GO terms, such as synaptic and sensory functions (Extended Data Fig. 6e). These neurons may also contribute to the atypical sociability observed in VPA-treated mice. However, we do not rule out the possibility that their transcriptomic alterations may be due to aberrant parvocellular PVH^OT^ neurons, as previous studies have reported that OT ligands are involved in the maturation of GABAergic neurons^31^.

In summary, our data suggest that the effects of embryonic VPA treatment on gene expression patterns are heterogeneous and cell type-specific. Certain gene expression networks relevant to critical neural functions are selectively vulnerable in some specific cell types within and near the PVH, including the parvocellular PVH^OT^ neurons.

### Activation of PVH^OT^ neurons to restore atypical social behavior

Chemogenetic activation^32^ of PVH^OT^ neurons has been widely employed to facilitate prosocial behaviors^16, 33^ and potentially ameliorate atypical sociability in rodent models, specifically in *Cntnap2*-deficient mice^34, 35^ and *Shank3*-deficient rats^36^. Similarly, intranasal or intraperitoneal administration of OT has demonstrated a positive impact on social behaviors in *Shank3b*-deficient mice^5^, *Maged1*-deficient mice^17^, inbred mouse strains (BALB/cByJ and C58/J)^37^, and VPA-treated rats^20^. However, the effects of such manipulations on the aberrant gene expression of PVH^OT^ neurons remain unknown. Given that intranasal or intraperitoneal OT administration may not directly activate endogenous OT neurons or central OT receptors^38^, we focused on the chemogenetic activation strategy. We first used an adeno-associated virus (AAV) vector to target hM3D-mCherry to the PVH^OT^ neurons of VPA-treated *OT-Cre* mice, with a control virus expressing only mCherry. Additionally, we prepared an hM3D-expressing control group that was embryonically exposed to saline (Fig. 4a). Social behaviors, as assessed using the three-chamber test at 5 weeks of age, were not affected by clozapine-N-oxide (CNO) administration in the saline-treated group (Fig. 4b). Atypical social behaviors were observed in VPA-treated mice expressing only hM3D without CNO or expressing mCherry with CNO administration. However, administration of a single dose of CNO to hM3D-expressing VPA-treated mice restored their atypical social behaviors (Fig. 4b, right). Thus, chemogenetic activation of PVH^OT^ neurons ameliorated VPA-induced social dysfunctions.

**Fig. 4:**
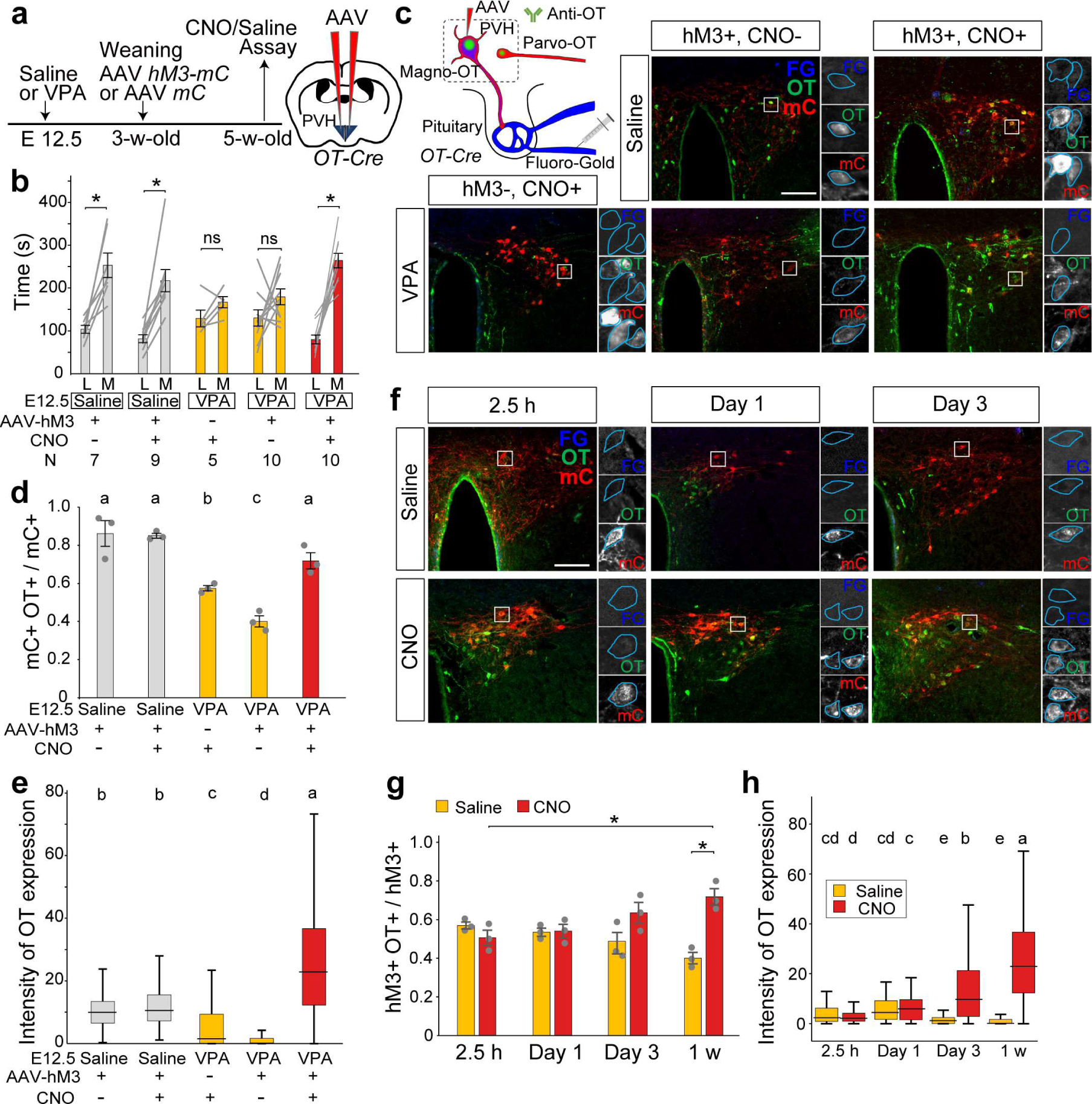
Chemogenetic activation of PVH^OT^ neurons rescues OT expression and atypical sociability. **a**, Experimental timeline and schematic of viral injection. mC, mCherry from AAVs. **b**, Duration of mice exploring a Lego block (L) and an unfamiliar male mouse (M). The number of animals (N) is indicated in the panel. *, p < 0.05 by the Wilcoxon rank-sum test with Bonferroni correction. **c**, Typical coronal sections containing parvocellular PVH^OT^ neurons (FG−) in the posterior parts of the PVH showing mCherry (hM3D or control) and anti-OT staining (OT+). Scale bar, 200 μm. **d**, **e**, Fraction of hM3D-mCherry+ OT+ cells over hM3D-mCherry+ cells (**d**) (N = 3 each) and fluorescent intensity of anti-OT staining in individual parvocellular PVH^OT^ neurons (**e**). Different letters (a–d) in the upper part of the graphs denote significant differences at p < 0.05 by one-way repeated measures ANOVA followed by the Tukey–Kramer post-hoc test (**d**) and one-way ANOVA with a post-hoc *t*-test with Bonferroni correction (**e**). **f**, Time course analysis of the fraction of hM3D-mCherry+ OT+ cells over hM3D-mCherry+ cells. **g**, Time course analyses of the fraction of hM3D-mCherry+ OT+ cells over hM3D-mCherry+ cells. *, p < 0.05 by the Wilcoxon rank-sum test. N = 3 mice each. **h**, Fluorescent intensity of anti-OT staining in individual parvocellular PVH^OT^ neurons. Different letters (a–e) in the upper part of the graph denote significant differences at p < 0.05 by two-way ANOVA with a post-hoc *t*-test with Bonferroni correction. Error bars, SEM.

Next, we generated hM3D-expressing VPA-treated or saline-treated control mice, which received a single dose of CNO 1 week before the histochemical analysis to quantify OT ligand expression. In the saline-treated control group, the administration of CNO did not have a significant impact on the level of OT ligand expression (Fig. 4c, d). By contrast, CNO administration significantly restored the reduced expression of OT ligands in the parvocellular PVH^OT^ neurons in the VPA-treated group (Fig. 4c–e). To assess the time course of this recovery, we analyzed brain samples obtained 2.5 hours, 1 day, 3 days, and 1 week after the single dose of CNO administration (Fig. 4f). We found that OT ligand expression in the parvocellular PVH^OT^ neurons was unaltered 2.5 hours or 1 day after CNO administration. Subsequently, OT ligand expression gradually and significantly increased (Fig. 4g, h). These results demonstrate that a single chemogenetic neurostimulation of PVH^OT^ neurons during the adolescent stage is sufficient to provide a prolonged rescue of abnormal OT ligand expression in parvocellular PVH^OT^ neurons.

Considering the prolonged effect observed, we hypothesized that the chemogenetic stimulation of OT neurons during the neonatal stage could potentially serve as a model of early intervention in atypical social behaviors. To investigate the consequence of prenatal VAP-exposure during the neonatal stage, we examined sections of the PVH from *OT-Cre*; *Ai9* double heterozygous male mice at postnatal day (PND) 2. We observed comparable numbers of tdTomato-positive cells in both saline- and VPA-treated mice, indicating the presence of Cre-mediated recombination in the PVH^OT^ neurons during the neonatal stage (Fig. 5a–c). In contrast to the data at 5 weeks of age (Fig. 1), the number of OT ligand-expressing cells remained unaltered in the VPA-treated group across the entire anterior–posterior axis of the PVH at PND 2 (Fig. 5a, b, d), suggesting that the reduction of OT expression in the parvocellular PVH^OT^ neurons occurs after PND 2.

**Fig. 5:**
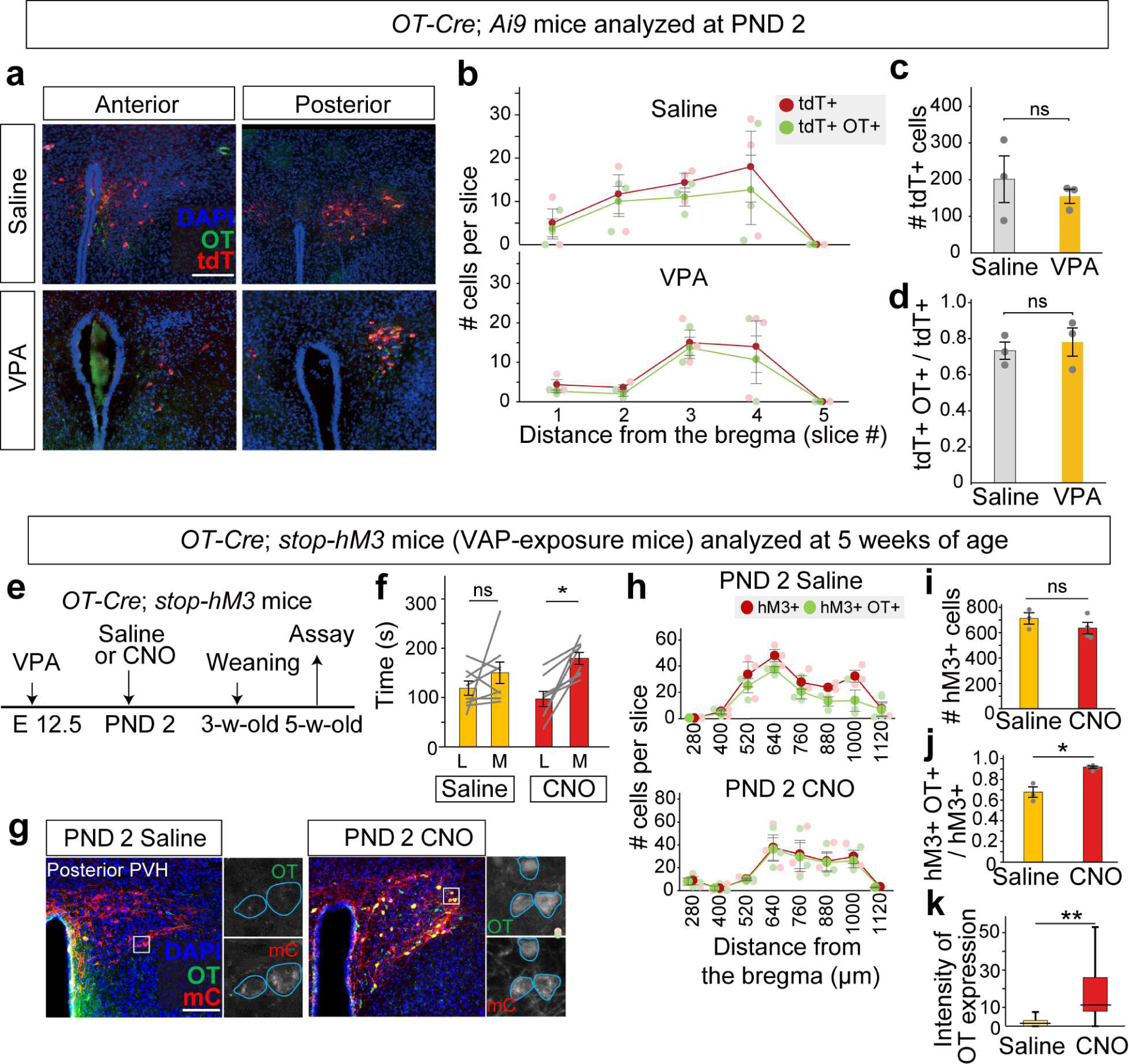
Neonatal activation of OT neurons persistently restores OT expression and social defects by VPA. **a**, Typical coronal sections of the anterior and posterior parts of the PVH from *OT-Cre*; *Ai9* double heterozygous male mice at PND 2 that prenatally received either saline (top) or VPA (bottom) injection. Scale bar, 200 μm. PND, postnatal day. **b**, Number of tdT+ cells and tdT+ OT+ dual-positive cells per slice along the anterior–posterior axis of the PVH. N = 3. **c**, Number of tdT+ cells, which is not significantly different between the saline and VPA groups by the Wilcoxon rank-sum test (N = 3 mice). **d**, Fraction of tdT+ OT+ cells over tdT+ cells. No difference was detected by the Wilcoxon rank-sum test (N = 3 mice). **e**, Timeline of the experiments. **f**, Duration of mice exploring a Lego block (L) and an unfamiliar male mouse (M). N = 8 each for the saline and CNO groups. *, p < 0.05 by the Wilcoxon rank-sum test. **g**, Typical coronal sections of the posterior PVH showing mCherry (inference of hM3D) and anti-OT staining (OT+). Scale bar, 200 μm. **h**, Number of hM3D-mCherry+ and hM3D-mCherry+ OT+ cells per slice along the anterior–posterior axis of the PVH. N = 3 for the saline and N = 4 for CNO groups. **i**, Number of hM3D-mCherry+ cells. No significant difference was detected by the Wilcoxon rank-sum test. **j**, The fraction of hM3D-mCherry+ OT+ cells over hM3D-mCherry+ cells. *, p < 0.05 by the Wilcoxon rank-sum test. **k**, Fluorescent intensity of anti-OT staining in individual cells as in Fig. 1g in the posterior PVH, where parvocellular PVH^OT^ neurons exist. **, p < 0.01 by the Wilcoxon rank-sum test.

To investigate the effects of chemogenetic activation of OT neurons at PND 2, we generated double heterozygous male mice by crossing *OT-Cre* mice with a mouse line that drives hM3D-mCherry in a Cre-dependent manner^39^. These mice were subjected to embryonic VPA treatment and received a dose of CNO on PND 2 (Fig. 5e). We confirmed the specificity of hM3D expression in these mice through histochemical analysis (Extended Data Fig. 7). Subsequently, we evaluated their social behaviors using the three-chamber test at 5 weeks of age. We observed that the CNO-treated group displayed improved sociability compared with the saline-treated control group (Fig. 5f). Although neonatal CNO administration did not alter the number of hM3D-expressing PVH^OT^ neurons, OT ligand expression in hM3D-expressing posterior PVH^OT^ neurons were significantly restored compared with the saline-treated control group (Fig. 5g–k). These findings indicate that targeted neurostimulation of OT neurons during the neonatal stage ameliorates both the OT ligand expression and sociability of VPA-treated mice during later young adulthood, providing a valuable model for early intervention in the atypical development of social behaviors.

### Transcriptomic analysis of PVH^OT^ neurons following chemogenetic stimulation

Next, we aimed to investigate the effects of neonatal OT neuron stimulation on gene expression in the PVH. We conducted snRNA-seq at 5 weeks of age using male *OT-Cre*; *stop-hM3* mice that had been exposed to VPA prenatally and treated with either CNO or saline as a control at PND 2 (Fig. 6a). We obtained snRNAseq profiles for 4635 and 4245 cells in the CNO- and saline-treated groups, respectively. Based on marker gene expression, we classified *vGluT2*+ clusters into one of the previously identified cell clusters based on Fig. 2b data (Fig. 6b, red, Extended Data Fig. 8a–d) and identified parvocellular and magnocellular PVH^OT^ neurons (Fig. 6c).

**Fig. 6:**
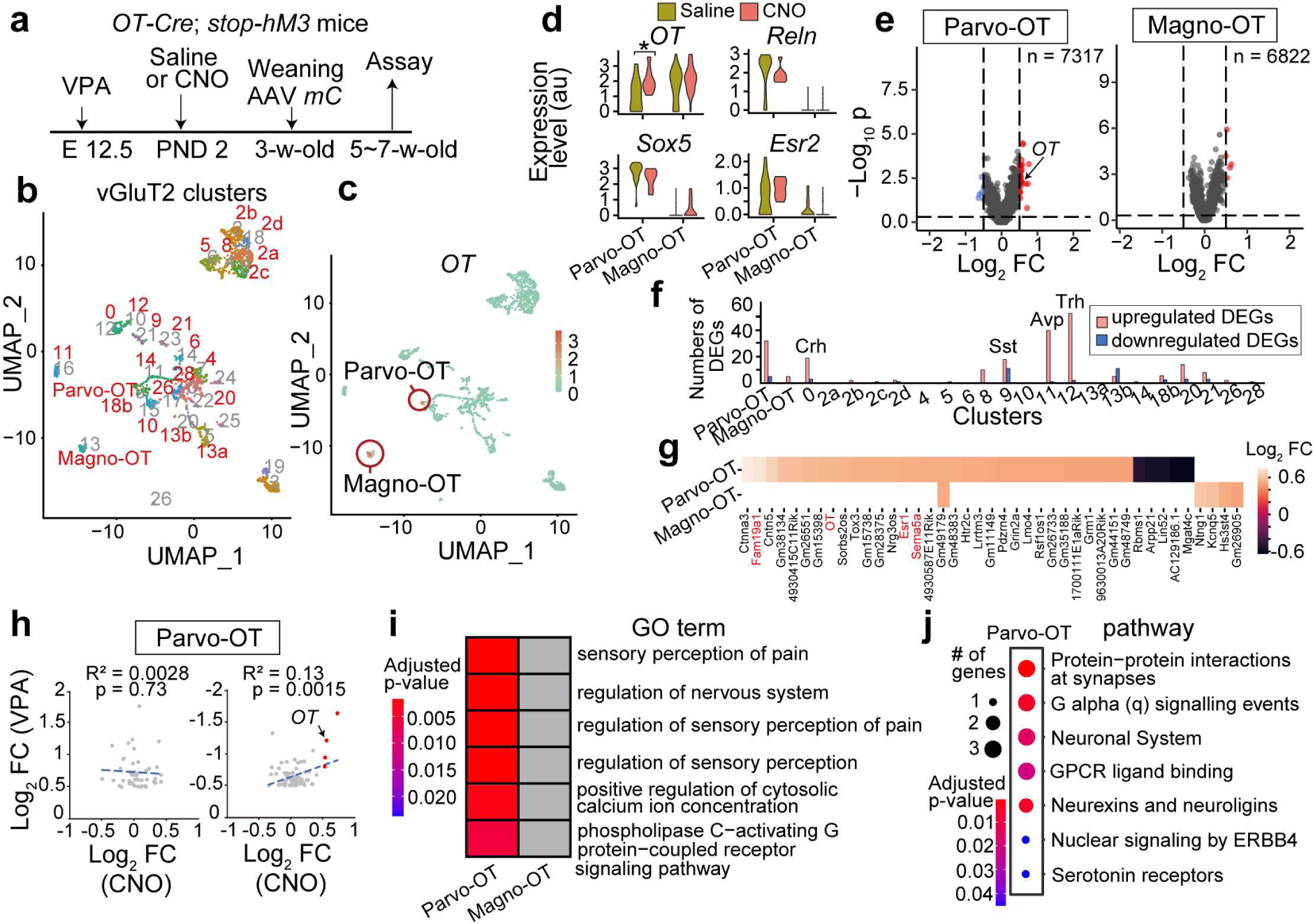
Long-lasting effects of neonatal stimulation of OT neurons. **a**, Timeline of the experiments. PND, postnatal day. **b**, **c**, UMAP of *vGluT2*+ cell clusters (**b**) and *OT* gene expression (**c**), with a color scale showing log-normalized expression. The clusters highlighted in red correspond to the *vGluT2*+ clusters in Fig. 2b based on marker gene expressions (Extended Data Fig. 8b). **d**, Violin plots showing *OT* gene expression, and three genetic markers (*Sox5*, *Reln*, and *Esr2* genes) used for distinguishing parvocellular and magnocellular PVH^OT^ neurons. *, p < 0.05 by the Wilcoxon rank-sum test. **e**, Volcano plots representing upregulated (red) and downregulated (blue) DEGs in parvocellular and magnocellular PVH^OT^ neurons, as in Fig. 2e. **f**, The numbers of upregulated (red) and downregulated (blue) DEGs within the *vGluT2+* clusters (cluster names are based on Fig. 2b). **g**, Heatmaps showing the log2-transformed fold change for DEGs found in parvocellular and magnocellular PVH^OT^ neurons. Genes downregulated in the VPA-treated group (Fig. 2e) are denoted in red. **h**, Correlation between the log2-transformed fold changes in the CNO-treated group (X-axis) and those in the VPA-treated group (Y-axis, left for upregulated and right for downregulated DEGs) as based on Fig. 2 data. Red dots represent significantly upregulated genes in the CNO-treated group. **i**, GO terms associated with the upregulated DEGs are shown with a heatmap of p-values for each GO term in parvocellular and magnocellular PVH^OT^ neurons. Grey cells indicate no enrichment. **j**, Heatmap representing p-values for pathway analysis using upregulated DEGs in the parvocellular PVH^OT^ neurons. For more data, See Extended Data Figs. 7–8 and Supplementary Table 4.

The DEG analysis unveiled more abundant upregulated genes within the CNO-treated group, particularly in parvocellular PVH^OT^ neurons (Fig. 6d–f, Supplementary Table 4). These DEGs showed minimal overlap between parvocellular and magnocellular PVH^OT^ neurons (Fig. 6g). Remarkably, the *OT* gene itself was significantly upregulated in parvocellular PVH^OT^ neurons of CNO-treated mice (Fig. 6d, e, g), indicating that the observed recovery in OT ligand expression (Fig. 5g–k) is a result of restored mRNA expression. Additionally, the downregulated DEGs in the parvocellular PVH^OT^ neurons of the VPA-treated group (Fig. 2) displayed a significant trend of increased expression in the CNO-treated group (Fig. 6h), indicating the recovery. In contrast, the upregulated DEGs in the VPA group did not show a trend of restoration. To further validate the gene expression recovery, we visualized the expression of some of the upregulated DEGs found in parvocellular PVH^OT^ neurons *in vivo* through ISH. All 4 genes that we examined, including the *OT* gene, exhibited a significant increase in mRNA expression (Extended Data Fig. 9). The GO and pathway analyses of the upregulated DEGs highlighted enriched functionally relevant GO terms and pathways in parvocellular PVH^OT^ neurons (Fig. 6i, j) but not in the magnocellular PVH^OT^ neurons (Extended Data Fig. 8e). Besides OT neurons, we observed a large number of upregulated DEGs in other *vGluT2*+ clusters, including AVP-expressing (cluster 11) and thyrotropin-releasing hormone-expressing (cluster 12) neurons (Fig. 6f), suggesting the presence of non-cell autonomous effects of neonatal OT stimulation on other cell types in the PVH (Extended Data Fig. 8f). In contrast to *vGluT2*+ clusters, we found only a minimal effect on the GABAergic neurons (Extended Data Fig. 8g, h).

In summary, these data demonstrate that neonatal OT neuron stimulation exerts a cell-type-specific and long-lasting positive influence on certain aberrant gene expressions in the PVH neurons, including parvocellular PVH^OT^ neurons, which potentially contributes to the restoration of social behaviors.

## DISCUSSION

Efforts to develop an effective treatment for atypical social traits in NDDs are impeded by our limited comprehension of the pathophysiology associated with the symptoms, particularly at the cellular level. Social dysfunction arises from a complex interplay between a multitude of genetic risk factors^40, 41^ and a wide range of maternal and environmental influences during fetal brain development^2, 3^, which have the potential to impact the entire nervous system. Previous research has predominantly centered on symptom-correlated traits in the cerebral cortex^42–44^, whereas the hypothalamic regulators of social behavior have received comparatively little attention^5, 12^. In the present study, we employ snRNAseq to elucidate the specific effects on the gene expression in the hypothalamic center regulating social behaviors, particularly focusing on social dysfunction and recovery models. Here, we discuss the biological insights yielded by our study.

Although diminished OT immunoreactivity in the PVH is common in both genetic^5, 16–18^ and environmental^4, 19, 20^ models of social dysfunctions, previous research has not distinguished cellular loss from a reduction of OT expression. Our data have ruled out cellular depletion in PVH^OT^ neurons in VPA-exposed mice. Instead, abnormalities in gene expression in parvocellular PVH^OT^ neurons, including the *OT* gene, are particularly concentrated in vital neural functions and signal transductions. Consequently, the ability of parvocellular PVH^OT^ neurons to convey information to downstream targets would be severely compromised. Analogous to the pronounced impacts on specific cell types seen in neurodegenerative disorders^1^, the framework of selective vulnerability at the level of gene expression can facilitate our understanding of social dysfunctions in a broader context. Future studies employing transcriptome analysis should explore whether the reduction in *OT* expression is common in various genetic social dysfunction models^5, 16–18^ and whether it stems from a shared vulnerability in the gene regulatory system within parvocellular PVH^OT^ neurons.

We have established that chemogenetic stimulation of OT neurons exerts sustained effects on gene regulation in the parvocellular PVH^OT^ neurons, correlating with the restoration of sociability. What might underlie these enduring effects? Given the absence of a decline in OT expression at the neonatal stage of VPA-treated mice (Fig. 5a–d), a plausible hypothesis is that prenatal factors, such as VPA exposure, could induce an epigenetic scar^19^, subsequently disrupting the crucial gene regulatory network for social behavioral development beyond infancy. Neonatal chemogenetic activation could mitigate these epigenetic scars, resulting in enduring effects. Future studies should characterize the epigenetic status at various stages of social behavioral development. In relation to this matter, it is important to comprehend the nature of stimulating OT neurons, encompassing an analysis of whether intense action potentials or specific signal transductions mediated by hM3D confer therapeutic benefits. Stimulating OT neurons with blood-brain barrier-permeable small molecular compounds may also be of interest, with potential relevance to application in humans. In this context, our snRNAseq data suggest that signaling pathways involving estrogen receptor ERα/β, serotonin receptor 5-HTR2c, and the PI3K/Akt could be plausible targets for stimulating parvocellular PVH^OT^ neurons (Figs. 3, 5 and Extended Data Fig. 2a). Collectively, our established model for early intervention, combined with snRNAseq analysis, provides a valuable platform for investigating effective treatment strategies and the underlying mechanisms.

Parvocellular PVH^OT^ neurons can modulate diverse brain functions^11, 15, 42^, including social reward^12, 14, 45^. Their local neural circuitry enables influence over magnocellular PVH^OT^ neurons^46^, thus exerting control over the entire OT system and collectively regulating various circuit functions^10^, such as the cortical excitatory/inhibitory balance^47, 48^. Additionally, parvocellular PVH^OT^ neurons impact gene expression across various cell types^31^, including AVP neurons (Fig. 6), which also contribute to social behaviors^9^. Targeted genetic manipulations of parvocellular PVH^OT^ neurons are crucial for dissecting their roles across diverse brain regions and cell types in regulating social behaviors. To achieve these goals in future investigations, utilizing a manipulation system based on the *OT* minipromoter^8, 15^, in conjunction with cell-type specific marker genes identified in this and the previous study^12^, holds promise.

## Methods

### Animals

All animal experiments were approved by the Institutional Animal Care and Use Committee of the RIKEN Kobe Branch (#A2017-15-13). *OT*-*Cre* mice (JAX #024234) were purchased from the Jackson Laboratory. *Rosa26^tm9(CAG–tdTomato)Hze^* mice (also known as *Ai9* mice) (JAX #007909) were kindly provided by Takeshi Imai, who purchased them from the Jackson Laboratory. *Rosa26^dreaddm3^* mice (also known as *stop-hM3* mice) were kindly provided by Takeshi Sakurai. All mice were maintained on a C57BL6 background. Animals were housed at the animal facility of the RIKEN Center for Biosystems Dynamics Research (BDR) under ambient temperature (18–23 °C) and a 12-h light, 12-h dark cycle schedule. The mice were allowed free access to the laboratory diet (MFG; Oriental Yeast, Shiga, Japan; 3.57 kcal/g) and water unless otherwise mentioned.

### VPA treatment

To generate a group of mice treated with VPA^19, 22^, we administrated sodium valproate (Sigma, #P4543) at a dosage of 500 mg/kg to pregnant female mice on gestation day 12.5, based on the identification of the mating day through the formation of a vaginal plug. For the control group of mice treated with saline, we injected 10 mL/kg saline to pregnant female mice at the same stage.

### Behavioral assays

The three-chamber test was performed using an apparatus consisting of three chambers measuring 20 × 30 × 30 cm (width × length × height), with the two side rooms connected to the central room by a small 5 × 3 cm (width × height) passageway. The test was conducted according to previously described methods^49^. Briefly, the animals were allowed to habituate to the empty chamber for 10 min. Then, the grids were placed in both side chambers, and the animals were further allowed to habituate for 10 min. A non-familiar C57BL/6 young adult mouse or a Lego block was placed on the grid and the animal’s behaviors were recorded for 10 min. The side chambers were cleaned with ethanol between each habituation and recording session. The recording was done using a GoPro8 camera (GoPro; #CHDHX-801-FW). The experiment was conducted approximately around zeitgeber time (ZT) 6, where ZT 0 was defined at the onset of the light period. For pharmacogenetic activation during adolescence (Fig. 4), behavior sessions were initiated 30 min after intraperitoneal administration of CNO (5 mg/kg, Tocris, #4936/10) or saline (200 μL). For pharmacogenetic activation during the neonatal stage (Fig. 5 and 6), CNO (5 mg/kg) or saline (10 μL) was orally administered on PND 2, and behavioral experiments were conducted at 5 weeks of age.

To evaluate social behaviors, we manually quantified the time spent by the mice exploring the grid containing the non-familiar mouse and the grid containing the object during the 10-min recording session.

### Single nucleus RNA sequencing (snRNA-seq): library preparation

The “Frankenstein” protocol (doi:10.17504/protocols.io.3fkgjkw) was modified to isolate the nucleus as follows. For Figs. 2–3 data, male *OT-Cre; Ai9* mice at 5 weeks of age were deeply anesthetized by isoflurane (Fujifilm, #099-06571), perfused by cold phosphate-buffered saline (PBS) to remove blood cells, and euthanized by decapitation. Brains were sectioned into 1000-μm coronal slices under microscopy and the sections were floated in 1% BSA-PBS (Nacalai Tesque, #0128197 and Takara, #T9181). The PVH was dissected based on the fluorescence of tdTomato in *OT-Cre; Ai9* male mice, and homogenized sufficiently in ice-cold Nuclei EZ Lysis Buffer (Millipore Sigma, N-3408). The resulting suspension was incubated on ice for 5 min and filtered through a 70-μm strainer. The nuclei were pelletized by centrifugation at 500×g and 4 °C for 5 min, and the supernatant was removed. The nuclei were resuspended in Nuclei EZ Lysis Buffer and pelletized again by centrifugation at 500×g and 4 °C for 5 min. They were then resuspended in 1% BSA-PBS containing 0.2 U/μL RNase Inhibitor (Roche, 3335399001) (Resuspension Buffer). We utilized 2–4 male mice for each condition, pooled the isolated nuclei, and centrifuged again at 500×g and 4 °C for 5 min to pellet the nuclei. The nuclei were then resuspended in resuspension buffer and 10 μg/mL DAPI, and filtered through a 40-μm strainer. The nuclei were sorted from the suspension using a cell sorter (SH800Z; Sony) at 5 °C. The gate was set first to identify a single nucleus population based on the DAPI signal and then to select larger nuclei preferentially, inferring neurons while simultaneously excluding multiplets (Extended Data Fig. 1a). Immediately after sorting was completed, the nuclei stained with DAPI were counted using a hemocytometer to verify the yield of intact nuclei. The nuclei were then resuspended in a resuspension buffer and used for generating snRNA libraries using the 10X Genomics Chromium platform targeting 8000 nuclei per condition. The libraries were prepared using the 10X Genomics RNA 3′ v3 kit (#1000269). The completed libraries were sequenced to a depth of 110 Gb on HiSeqX (performed by Novogene).

For Fig. 6 data, we utilized male *OT-Cre; stop-hM3* mice at 5–7 weeks of age that had been exposed to VPA prenatally and CNO or saline as a control at PND 2. We followed the above protocols with slight modifications. Briefly, *OT-Cre; stop-hM3* mice were injected with AAV in the PVH and the supraoptic nucleus regions at three weeks of age to facilitate visualization of OT neurons during the dissection process. To optimize the yield in collecting neural cell nuclei, the FACS gating was adjusted to accommodate larger nuclei. Each batch aimed to isolate 5,000 nuclei using the Chromium system. The completed libraries were sequenced to a depth of 105 Gb on HiSeqX (performed by AZENTA).

### snRNA-seq: data analysis

To conduct the analysis, we aligned the fastq files from each library to the mm10 reference transcriptome (mm10, gencode version vm23) using the Cell Ranger pipeline. After alignment, we loaded feature barcode matrices into Seurat (R package, v4.3.0). Cells were retained if at least 800 Unique Molecular Identifiers (UMIs) were detected, and genes were retained if at least one UMI was detected in at least three cells. In Figs. 2–3 data, after quality control and initial filtering, we recovered 5491 and 10,904 total nuclei from the saline and VPA libraries, respectively. The median numbers of genes/nuclei were 3232 for the saline library and 2460 for the VPA library. The saline library contained a considerable number of cells that appeared to be doublets, as evidenced by having more than twice the mean number of UMI and barcodes. To address this issue, we utilized Cloupe to confirm that a cell population containing 4300 or more barcodes formed a doublet cluster, and then removed such cells. Similarly, we removed cells with more than twice the mean number of UMI in the VPA library. The saline and VPA libraries were integrated using the sctransform function in Seurat. Normalization of the count data was conducted using LogNormalize. In this method, the following calculations were made. Feature counts for each cell were divided by the total counts for that cell and multiplied by the scale factor. This was then natural-log transformed using log1p.

We identified the top 3000 variable features and employed them as input for principal component analysis. We performed the initial clustering (Extended Data Fig. 1) using the top 35 principal components (PCs) at a resolution of 0.02 to distinguish between neuronal and non-neuronal cell populations. Based on the expression of *Camk2a*, *Slc17a6*, *Gad1*, and *Gad2*, we identified the neuronal cell population and utilized the subsetUmapClust function to perform re-clustering. Furthermore, we separated the neuronal population into vGluT2+ and Gad1/2+ subpopulations based on the expression of *Slc17a6*, *Gad1*, and *Gad2* using the same parameters as before. We identified 1133 and 3225 nuclei in the saline and VPA libraries for the vGluT2+ neuronal population, respectively, which collectively constituted 30 clusters using the top 40 PCs at a resolution of 1.00. In Fig. 2a and b, we counted as follows. In the saline library, 17 and 21 nuclei were classified as magnocellular and parvocellular OT neurons, respectively, and in the VPA library, 94 and 90 nuclei were classified as magnocellular and parvocellular OT neurons, respectively. In Extended Data Fig. 6, we identified 17 clusters in the GAD1/2 neuronal population using the top 40 PCs at a resolution of 0.25.

For Fig. 6 data, we followed the aforementioned procedures with minor adjustments. Briefly, we collected two batches, labeled as 1 and 2, for the CNO group, while data for the saline group were derived from a single batch. For both saline and CNO batch 1, we applied the method outlined above to eliminate nuclei forming doublets. Consequently, we removed nuclei with more than 5500 barcodes for saline and 5800 barcodes for CNO batch 1. Notably, in CNO batch 2, a considerable number of nuclei exhibited a trend of RNA degradation, as indicated by the Cell Ranger barcode rank plot. As a result, we focused our analysis on nuclei containing over 4000 barcodes based on the observation that parvocellular PVH^OT^ neurons found in saline and CNO batch 1 exhibited this range of barcodes per cell. We set the upper limit for barcodes in CNO batch 2 at 7000, considering that cells with barcodes approximately 1.8 times the lower limit could be singular. We performed the initial clustering using the top 50 PCs at a resolution of 0.02 to distinguish between neuronal and non-neuronal cell populations. Based on the expression of *Camk2a*, *Slc17a6*, *Gad1*, and *Gad2*, we identified the neuronal cell population and utilized the subsetUmapClust function to perform re-clustering using the top 30 PCs at a resolution of 0.50. In the saline and CNO libraries, we identified 880 and 1242 nuclei, respectively, within the vGluT2+ neuronal population, constituting a total of 27 clusters using the top 50 PCs at a resolution of 1.50.

In Extended Data Fig. 8, we identified magnocellular and parvocellular PVH^OT^ neurons in cluster 13 (corresponding to cluster 19 in Fig. 2) and cluster 9 (corresponding to cluster 18 in Fig. 2) within the vGluT2+ neuronal population using the top 20 PCs at a resolution of 0.9. In Fig. 6 and Extended Data Fig. 8, we counted as follows: In the saline library, 35 and 19 nuclei were classified as magnocellular and parvocellular OT neurons, respectively, while in the CNO library, 37 and 12 nuclei were classified as magnocellular and parvocellular OT neurons, respectively. The violin plots (Fig. 6d) for the CNO group were based on data from a representative batch.

Regarding the GAD1/2+ clusters in Extended Data Fig. 8, we limited our comparison to the saline and CNO batch 1, as CNO batch 2 had a lower number of barcodes. Utilizing the top 40 PCs at a resolution of 0.5, we identified 20 clusters and analyzed 1558 and 1348 nuclei in the saline and CNO libraries for the GAD1/2+ neuronal population, respectively.

### snRNA-seq: DEGs, GO, and pathway analysis

We considered a gene to be “expressed” in a given cell type if at least one UMI was detected in 30% or more of the cells of that type. We then identified DEGs that exhibited upregulation or downregulation under the specific comparison conditions. DEGs were defined by an absolute log2-fold change (|log2-FC|) greater than 0.5 and a p-value less than 0.05. A curated list of highly confident autism-associated genes was obtained from the SFARI Gene database (https://gene.sfari.org) as of 26 September 2023. In Fig. 2 and Supplementary Table 1, “SYNDROMIC”, “CATEGORY 1”, “CATEGORY 2”, “CATEGORY 3” and “Archive only” genes in the SFARI Gene database were referred to as “S”, “1”, “2”, “3”, and “Acv”, respectively.

In Figs. 3, 6, Extended Data Figs. 6, 8, and Supplementary Tables 2 and 3, we conducted GO and pathway analyses of biological processes using clusterProfiler^27–29^ for each cell cluster based on DEGs that were up- or down-regulated under the specified comparison conditions. The GO terms and pathways were sourced from an established compendium of designated terminologies. In cases where a single gene mapped to multiple isoform pathways, these were consolidated and treated as a singular pathway for the analysis.

### Maternal high-fat diet (MHFD)

Female mice were housed with either a regular diet (RD) consisting of 5.5% of total kilocalories (kcal) from fat, 27.8% kcal from protein, and 48.6% kcal from carbohydrates (MFG; Oriental Yeast; 3.57 kcal/g), or a high-fat diet (HFD) consisting of 62.2% kcal from fat, 18.2% kcal from protein, and 19.6% kcal from carbohydrates (HFD-60; Oriental Yeast). After 8 weeks on the respective diets, female mice were mated with *OT-Cre* adult males. The HFD was continued throughout the pregnancy and lactation periods. The resultant offspring were weaned at 3 weeks of age and all were transitioned to the RD, regardless of their mothers’ dietary condition.

### Viral preparations

The following AAV vectors were generated by the Gunma University Viral Vector Core and Addgene using the corresponding plasmids (Addgene #44361, #27056, and #184754).

AAV serotype 8 *hSyn-DIO-hM3Dq-mCherry* (2.1 × 10^13^ gp/mL)

AAV serotype 8 *hSyn-DIO-mCherry* (2.3 × 10^13^ gp/mL)

AAV serotype 9 *OTp-mCherry* (2.3 × 10^13^ gp/mL)

### Stereotactic and orbital venous plexus injections

Male mice (*OT-Cre*, *OT-Cre; Ai9*, and *OT-Cre; stop-hM3*) at the age of 3 weeks were used for injection. All stereotaxic injections were performed under general ketamine–xylazine anesthesia, with 65 mg/kg ketamine (Daiichi-Sankyo) and 15 mg/kg xylazine (Sigma-Aldrich, cat# X1251), using a stereotaxic instrument (RWD, cat#68045). The coordinates for injection into the PVH were as follows: anterior 0.78 mm and lateral ± 0.1 mm from the bregma, and ventral 4.5 mm from the brain surface. 250 nL of either AAV8 *hSyn-DIO-hM3Dq-mCherry* or AAV5 *hSyn-DIO-mCherry* was bilaterally injected into the PVH at a speed of 50 nL/min using a UMP3 pump regulated by Micro-4 (World Precision Instruments). The coordinates for injection into the supraoptic nucleus were as follows: anterior 0.78 mm and lateral ± 1.2 mm from the bregma, and ventral 5.5 mm from the brain surface. 250 nL of either AAV serotype 9 *OTp-mCherry* was bilaterally injected into the PVH and the supraoptic nucleus at a speed of 50 nL/min using a UMP3 pump regulated by Micro-4 (World Precision Instruments).

To achieve selective labeling of magnocellular PVH^OT^ neurons with FG, 30 μL of 2% FG (Fluorochrome, cat#526-94003) in saline was administered either unilaterally or bilaterally into the orbital venous plexus using a 1-mL syringe and 25-G needle. Animals were sacrificed for histochemical analysis at least 24 h after FG injection.

### Histology and histochemistry

Brains from *OT-Cre*, *OT-Cre; Ai9*, and *OT-Cre; stop-hM3* mice were subjected to immunolabeling. The mice were anesthetized with an overdose of isoflurane and then perfused transcardially with PBS, followed by 4% paraformaldehyde (PFA) in PBS. Brain tissues were post-fixed overnight with 4% PFA in PBS at 4 °C, cryoprotected with 30% sucrose solution in PBS for at least 24 h, and embedded in O.C.T. compound (Tissue-Tek, cat#4583). We collected 30-μm coronal sections of the whole brain using a cryostat (model #CM1860; Leica) and placed them on MAS-coated glass slides (Matsunami, cat#MAS-13). The following primary antisera were used for immunolabeling: rabbit anti-OT (1:500, IMMUNOSTAR, cat#20068) and goat anti-mCherry (1:500, Acris Antibodies GmbH, cat# ACR-AB0040-200-0.6). The primary antisera were detected with the following secondary antibodies: donkey anti-rabbit Alexa Fluor 488 (1:250, Invitrogen, cat# A32790), donkey anti-goat Alexa Fluor 555 (1:500, Invitrogen, cat# A32816), and donkey anti-rabbit Alexa Fluor 647 (1:250, Invitrogen, cat#A31573). Sections were counterstained with DAPI (2.5 μg/mL) and imaged using an Olympus BX53 microscope equipped with a 10× objective lens (numerical aperture 0.4).

### *In situ* hybridization (ISH)

Fluorescent ISH was performed as previously described^50^. In brief, mice were deeply anesthetized with isoflurane and perfused with PBS, followed by 4% PFA in PBS. The brain was post-fixed with 4% PFA overnight. 30-μm coronal brain sections were made using a cryostat (Leica). To generate cRNA probes, DNA templates were amplified by PCR from the C57BL/6j mouse genome or whole-brain cDNA (Genostaff, cat#MD-01). T3 RNA polymerase recognition site (5’-AATTAACCCTCACTAAAGGG) was added to the 3’ end of the reverse primers. Primer sets to generate DNA templates for cRNA probes are as follows (the first one, forward primer, the second one, reverse primer):

*Esr2*-1 5’- AGACAAGAACCGGCGTAAAA; 5’- CGTGTGAGCATTCAGCATCT

*Esr2*-2 5’- CTCAATCTCGGGGTCTGAGT; 5’- CCAAGCAGGAAGAAAGAGGA

*Tsc1*-1 5’- AGGGGTTGCCTCACCTTACT; 5’- GGCACAGTCCTCCAACCTTA

*Tsc1*-2 5’- GGGTGGAGGCTTCTGTTTTA; 5’- ACCAGGGTGTCCTGTGTCTC

*Tsc1*-3 5’- CTCAGTTTGCACTGCAGCAT; 5’- TGGCATCTGATTTGGATTGT

*Sox5*-1 5’- CCCTCCATGTGGGAATAGAA; 5’- CAAACTTGAGGGTGGCATTT

*Sox5*-2 5’- AACGCATACAATGTGAAAACAGA; 5’- GGGCTTTAACCCATTTTCTTC

*Sema5a*-1 5’- CCCAACAAATCAAGCCAAAT; 5’- ATTTGCACCAGGCTCTGAAT

*Sema5a*-2 5’- CTCCACCCCTTCATAATCCA; 5’- TGGTGCATCTTATTGGCAGA

*Sema5a*-3 5’- TGACTTGACACCTGCTACGC; 5’- TGTTTCTTAAATGGCAGGGTTT

*Prickle1*-1 5’- GAGGACCGCAGCTCTCAAC; 5’- ACCGAGGCTTGAGCAGTTC

*Prickle1*-2 5’- CGCTAACGAGGAATTCTGGA; 5’- AGTCTGTCCTTGGCGGAGTA

*Prickle1*-3 5’- GACTCGTGGTGTTCGTCCTC; 5’- ACAAGCAGTCACCTCATCCA

*Ntrk2*-1 5’- GACTGAGCCTGGAGATTTGC; 5’- CAAACCTGGAATGGAATGCT

*Ntrk2*-2 5’- TGCACACATGTAGTGTGTTTGTG; 5’- AGTGATGAATCCCTCCCAAC

*Ntrk2*-3 5’- ATTTTGCAGCCTACGCATTC; 5’- AGGGTGAGAGAAGCTGGTCA

*Erbb4*-1 5’- TGATGAGGACAATGACAAATGA; 5’- AGCATTCCAAAGGTGCTGAC

*Erbb4*-2 5’- TGGGGCAAATAGGAAATTGT; 5’- GCATTTGAAGGCAAAGGCTA

*Erbb4*-3 5’- CCTCCTGTGACTTTTGTTGGA; 5’- GTGCATGTGCCATGAATGAT

*OT* 5’-TGGCTTACTGGCTCTGACCT; 5’- AGGAAGCGCGCTAAAGGTAT

*Esr1*-1 5’- TAAGAAGAATAGCCCTGCCTTG; 5’- ACAGTGTACGCAGGAGACAGAA

*Esr1*-2 5’- AGGCATGGTGGAGATCTTTG; 5’- AAGCCATGAGATCGCTTTGT

*Fam19a1*-1 5’- GCATTCATTTGGGGATTCAC; 5’- GCCAGAACGAGTTTCAGAGG

*Fam19a1*-2 5’- GCTACTGAATGCCTGGGAAA; 5’- AAGAGATCCACTTGGCTTGC

*Fam19a1*-3 5’- TGTGAGGTGGCTGGTGTATC; 5’- TCAGAGTGACCCACATGGAA

*Htr2c*-1 5’- GGTGCACCAGGCTTAATGAT; 5’- GAGACAGGGGCATGACAAGT

*Htr2c*-2 5’- ATGCACATGACTGTGGTGGT; 5’- AGCAGGTCCACGAATGAAAC

*Htr2c*-3 5’- CAGCTACTTGCACACCTTGG; 5’- GCAGTCTGTTGCACGTGTCT

DNA templates (500–1000 ng) amplified by PCR were subjected to in vitro transcription with DIG (cat#11277073910) or Flu (cat#11685619910)-RNA labeling mix and T3 RNA polymerase (cat#11031163001) according to the manufacturer’s instructions (Roche Applied Science). When possible, up to three independent RNA probes were mixed to increase the signal/noise ratio. For ISH combined with anti-mCherry staining, after hybridization and washing, sections were incubated with horseradish peroxidase-conjugated anti-Dig (Roche Applied Science cat#11207733910, 1:500) and goat anti-mCherry (1:500, Acris Antibodies GmbH, cat# ACR-AB0040-200-0.6) antibodies overnight. Signals were amplified by TSA-plus Biotin (AKOYA Bioscience, NEL749A001KT, 1:70 in 1× plus amplification diluent) for 25 min, followed by washing, and then mCherry-positive cells were visualized by donkey anti-goat Alexa Fluor 555 (1:500, Invitrogen, cat# A32816).

### Quantification and statistics: snRNA-seq data

All statistical analyses for sequencing data were performed in RStudio and R. VPA data were performed in RStudio Server v1.4.1717 and R v2.1.2. CNO data were performed in RStudio Server 2023.06.1 Build 524 and R v4.3.1. GO term and pathway analyses were performed in RStudio 2022.12.0+353 and R v4.2.2. Statistical tests and criteria used for snRNA-seq analyses are described in the relevant method details sections. The number of biological replicates for each experiment is stated in the relevant figure legends.

### Quantification and statistics: Histochemistry and ISH data

All image analysis was performed using napari software (doi:10.5281/zenodo.3555620, napari version 0.4.15, Python version 3.8.0, and Numpy version 1.23.1). To assess the intensity of anti-OT immunostaining and RNA probes in individual parvocellular PVH^OT^ neurons, we selected coronal sections located posteriorly at 1000–1120 μm from the bregma. Similarly, for magnocellular PVH^OT^ neurons, we opted for coronal sections located posteriorly at 520–640 μm from the bregma. By utilizing the labels tool in napari, all tdTomato- or hM3-mCherry-positive cells present within these sections were selected as regions of interest (ROIs), and the fluorescence intensity at each ROI was calculated using napari-skimage-regionprops (version 0.5.3). We randomly collected 100–300 ROIs for OT+ cells from at least three animals for each condition. Finally, the background fluorescence intensity was subtracted, and statistical analysis was conducted utilizing the corrected fluorescent intensity. Of note, the scales in Fig. 1g and 1n are different as the secondary antibodies used to detect anti-OT were different in these experiments because of a technical issue.

### Statistics

Statistical tests were performed using Excel, Python, R, and js-STAR (version 1.6.0). All tests were two-tailed. The sample size and statistical tests used are indicated in the figure legends. Statistical significance was set at p < 0.05 unless otherwise mentioned. Mean ± standard error of the mean (SEM) was used to report statistics unless otherwise indicated. In the box-and-whisker plots, the horizontal line within the box denotes the median, while the upper and lower sides of the box symbolize the first quartile and the third quartile, respectively. Two whiskers from the upper and lower edges, respectively, encompass the maximum and minimum values within a distance that extends to 1.5 times the interquartile range. Any outliers beyond this range are excluded from the plot.

## Other Supplementary Materials

**Supplementary Table 1: DEGs and ASD risk factor DEGs within the vGluT2+ clusters in VPA-exposed mice, related to Fig. 2**. This table provides the list of all DEGs with the p-value and log2FC value within the top 25 vGluT2+ clusters. We also manually identified DEGs that were designated as high-confidence ASD risk factor genes within each *vGluT2*+ cluster and shown in colored cells. The rank of ASD risk is also based on ref. 21 and https://gene.sfari.org/ as of 26 September 2023.

**Supplementary Table 2: GO terms that satisfy a false discovery rate criterion of < 0.01 in each *vGluT2*+ or *GAD2*+ cluster of VPA-exposed mice, related to Fig. 3**.

Supplementary Table 3: Pathways that satisfy a false discovery rate criterion of < 0.01 in each *vGluT2*+ cluster of VPA-exposed mice, related to Fig. 3.

**Supplementary Table 4: DEGs within the vGluT2+ clusters in CNO-mediated recovery experiments, related to Fig. 6**. This table provides the list of all DEGs with the p-value and log2FC value within the designated vGluT2+ clusters.

## Acknowledgments

We thank Kota Tamada and Toru Takumi (Kobe University), Teruhiro Okuyama (the Univ. of Tokyo) and members of the Miyamichi Lab for the critical reading of the manuscript, Ritsuko Morita and Hironobu Fujiwara for sharing the Sony SH800 cell sorter, Quan Wu and the DNA Analysis Facility at the Laboratory for Phyloinformatics for the advice on the snRNA-seq analysis, Masato Kinoshita and Hideki Enomoto for the advice on the MHFD, Takeshi Sakurai for sharing *stop-hM3* mice, Addgene for the AAV productions, and the animal facility of RIKEN BDR for taking care of the animals. AAV *OTp-mCherry* was produced by the Viral Vector Core of Gunma University Initiative for Advanced Research (GIAR) under the support of the Brain/MINDS program from AMED JP20dm0207057 and JP21dm0207111 to Hirokazu Hirai. This work was supported by a RIKEN Junior Research Associate Program to M.T., JSPS KAKENHI (20K20589, 21H02587), and a RIKEN Center Project Grant to K.M.

## Author contributions

M.T. and K.M. conceived the experiments. M.T. performed all the experiments and analyzed the data, with technical support from T.G., M.H., and S.I. M.T. and K.M. wrote the paper with help from all the authors.

## Declaration of interests

The authors declare that they have no competing interests.

## Data and materials availability

snRNaseq data have been deposited at Gene Expression Omnibus and are publicly available upon publication (GEO: GSE245555). This paper does not report the original code. All Python and R scripts used in this manuscript are available from the corresponding author upon reasonable request. Any additional information required to reanalyze the data reported in this paper is available from the corresponding author upon reasonable request.

---

**Extended Data Fig. 1:**
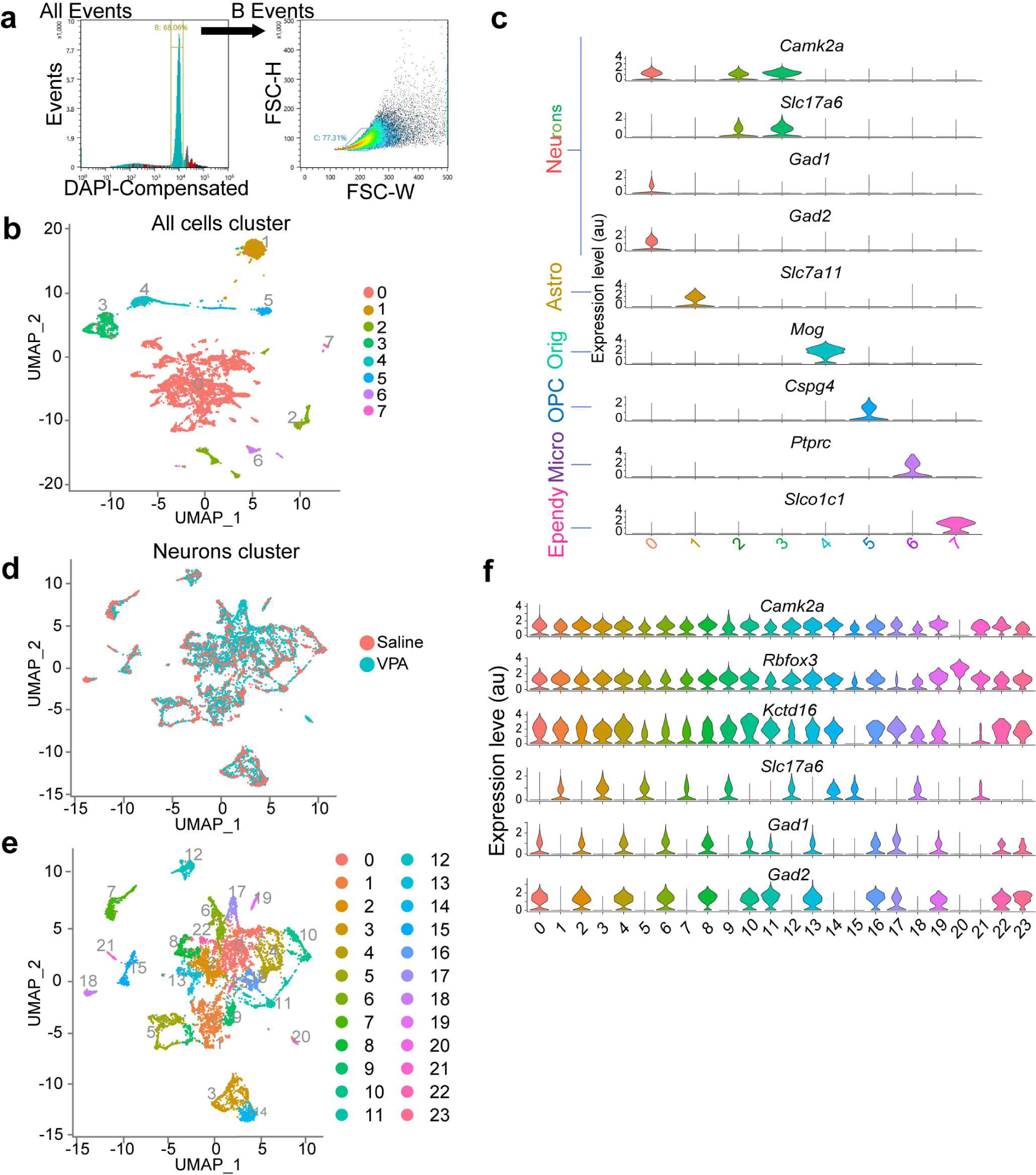
Global classification of snRNA-seq data, related to Fig. 2. **a**, Representative FACS gating to select DAPI-positive singlet nuclei (Gate B), followed by the exclusion of multiplets, while simultaneously choosing larger nuclei (inference of neurons) through FSC-W and SSC-H gating (Gate C). **b**, UMAP of all cells analyzed in this study showing seven clusters. **c**, Violin plots of indicated marker genes of various cell types. Based on their expression patterns, we annotated clusters 0, 2, and 3 as neurons, cluster 1 as astrocytes, cluster 4 as oligodendrocytes, cluster 5 as oligodendrocyte progenitor cells (OPCs), cluster 6 as microglia, and cluster 7 as ependymal cells. **d**, UMAP representation of the saline and VPA groups among the neuron cluster (a combination of clusters 0, 2, and 3 in panel **b**). **e**, UMAP representations of 24 clusters within the neuron cluster. **f**, Violin plots of indicated neural marker genes. We annotated clusters 1, 3, 5, 7, 9, 12, 14, 15, 18, and 21 as excitatory neurons based on the expression of the *vGluT2* (also known as *Slc17a6*) gene, and the rest as inhibitory neurons expressing *GAD1/2*.

**Extended Data Fig. 2:**
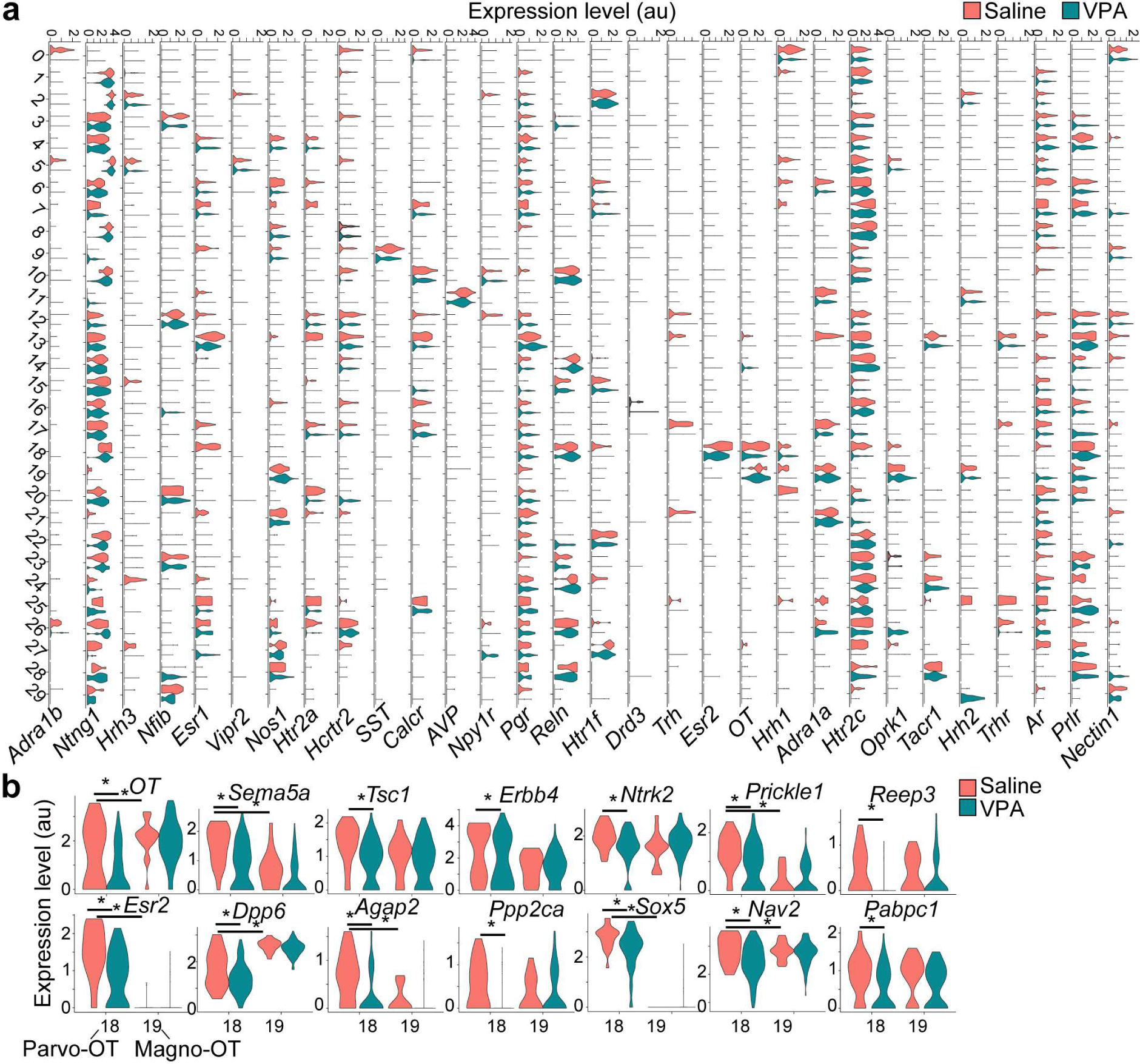
Clusters of *vGluT2+* neurons and the downregulated DEGs in the parvocellular PVH^OT^ neurons, related to Fig. 2. **a**, Violin plots displaying the expression of selected marker genes in the 30 clusters identified within the *vGluT2*+ neuron cluster (Fig. 1e), constructed based on Extended Data Fig. 1f. **b**, Violin plots illustrating the expression of downregulated DEGs in the parvocellular (cluster 18) and magnocellular (cluster 19) PVH^OT^ neurons. *, p < .05 by the Wilcoxon rank-sum test.

**Extended Data Fig. 3:**
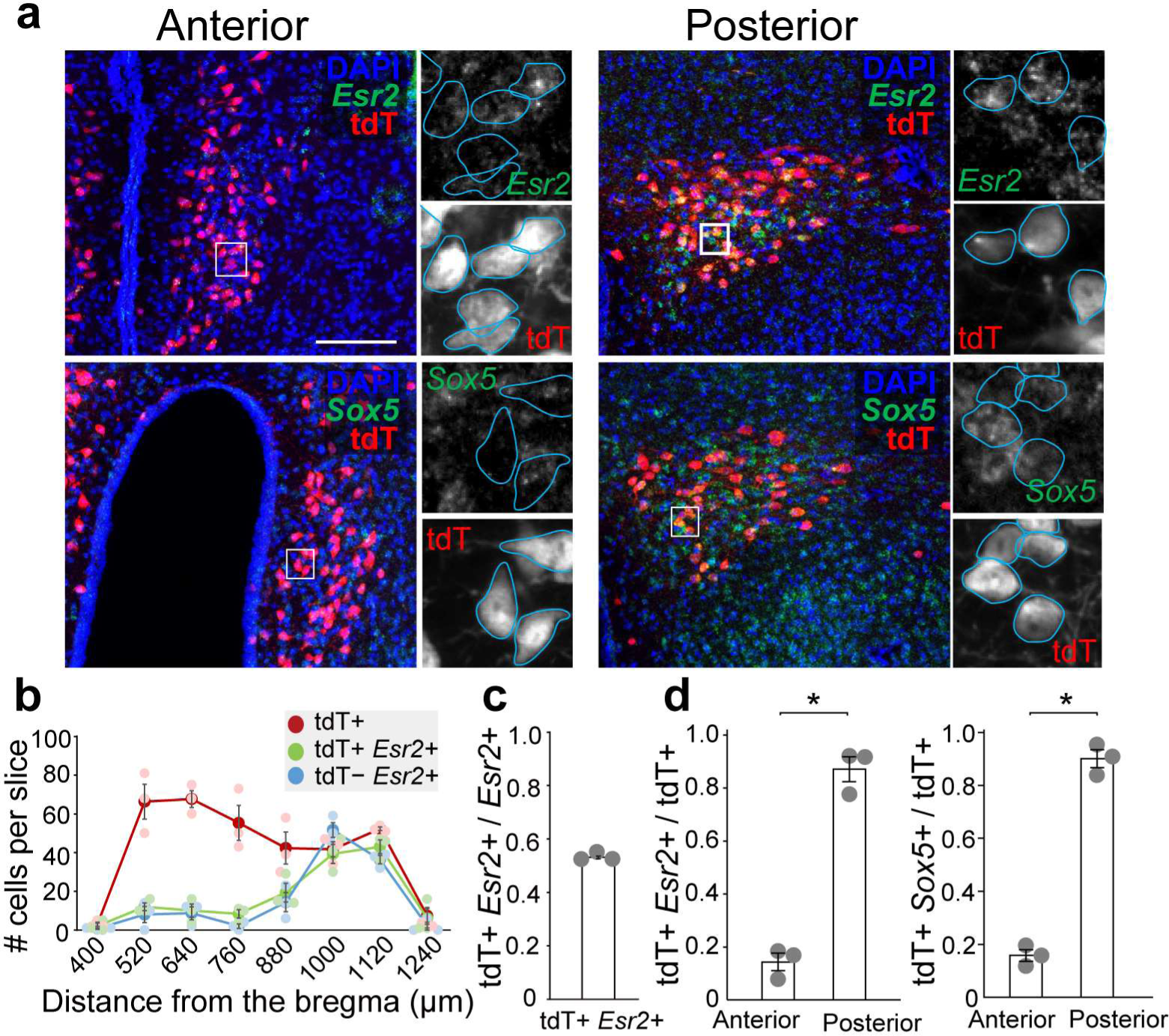
Expression patterns of marker genes for parvocellular PVH^OT^ neurons along the anterior–posterior axis, related to Fig. 2. **a**, Typical coronal sections of the anterior and posterior parts of the PVH from typically-developed (saline-injected) control *OT-Cre*; *Ai9* mice showing tdTomato (tdT) and RNA expressions for indicated genes. Scale bar, 200 μm. **b**, Number of tdT+, tdT+ *Esr2*+, and tdT− *Esr2*+ cells per slice along the anterior–posterior axis of the PVH. Error bars, SEM. N = 3 mice. **c**, Fraction of tdT+ *Esr2*+ cells over *Esr2*+ cells. **d**, Fraction of dually labeled cells over tdT+ cells for *Esr2* (left) and *Sox5* (right). Anterior and posterior sections correspond to posterior 640–760 μm and 1000–1120 μm from the bregma along the anterior–posterior axis. N = 3 each. *, p < 0.05 by the Wilcoxon rank-sum test. Error bars, SEM. Of note, the distributions of these marker genes resemble the cellular distribution of parvocellular PVH^OT^ neurons^12^.

**Extended Data Fig. 4:**
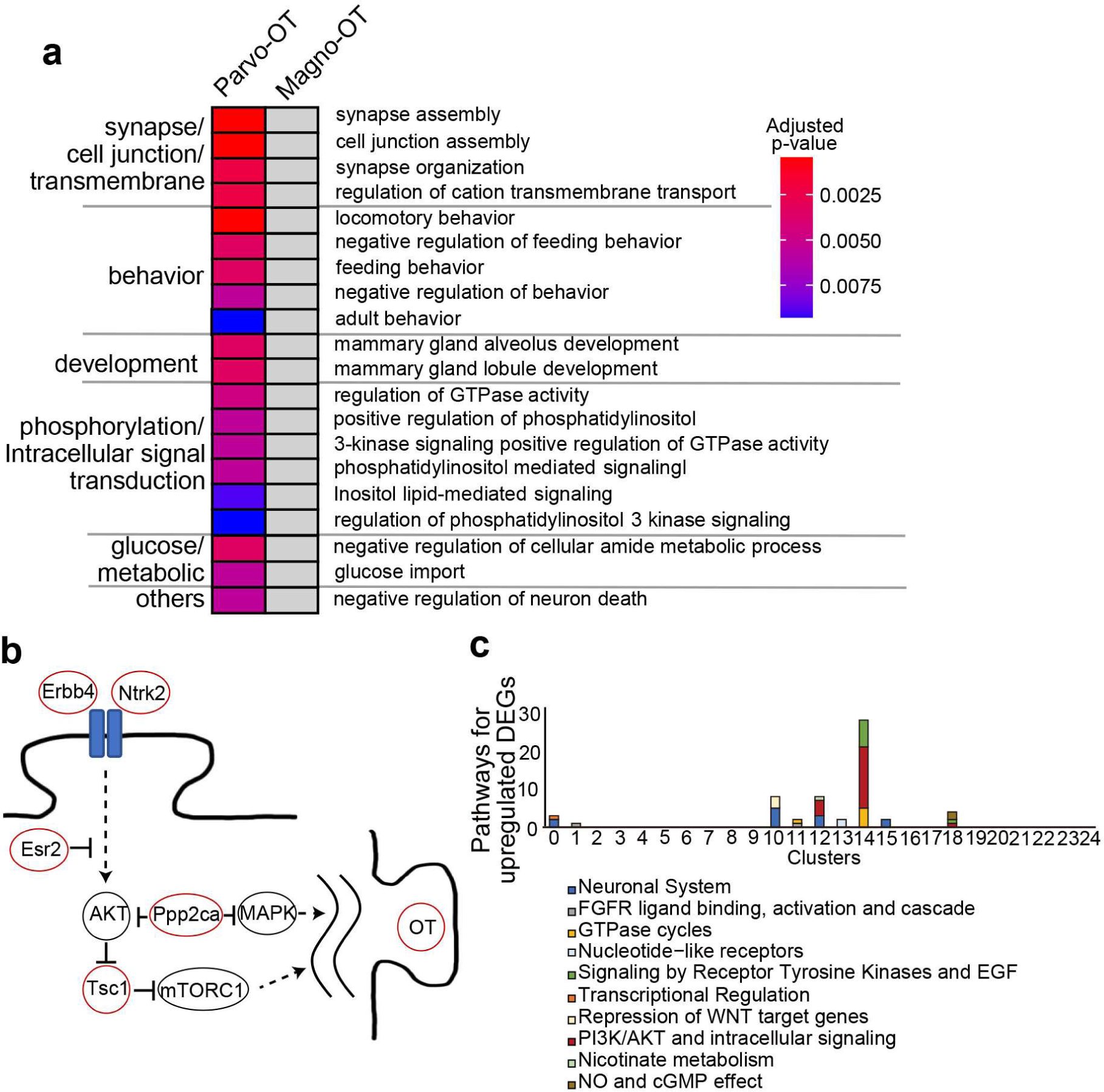
Gene Ontology (GO) and pathway analyses of PVH^OT^ neurons, related to Fig. 3. **a**, Downregulated DEGs were subjected to GO analysis. Specific GO terms are shown with a heatmap of p-values for each GO term in parvocellular and magnocellular PVH^OT^ neurons. Grey cells indicate no enrichment. **b**, Schematic diagram showing putative signaling pathways affected in the parvocellular PVH^OT^ neurons based on the downregulated DEGs, with red circles highlighting the products of ASD risk factor DEGs. These pathways are depicted based on the previous literature^30, 40, 51, 52^. **c**, The number of pathways that show significant association (p < 0.05) with upregulated DEGs within the top 25 *vGluT2+* clusters.

**Extended Data Fig. 5:**
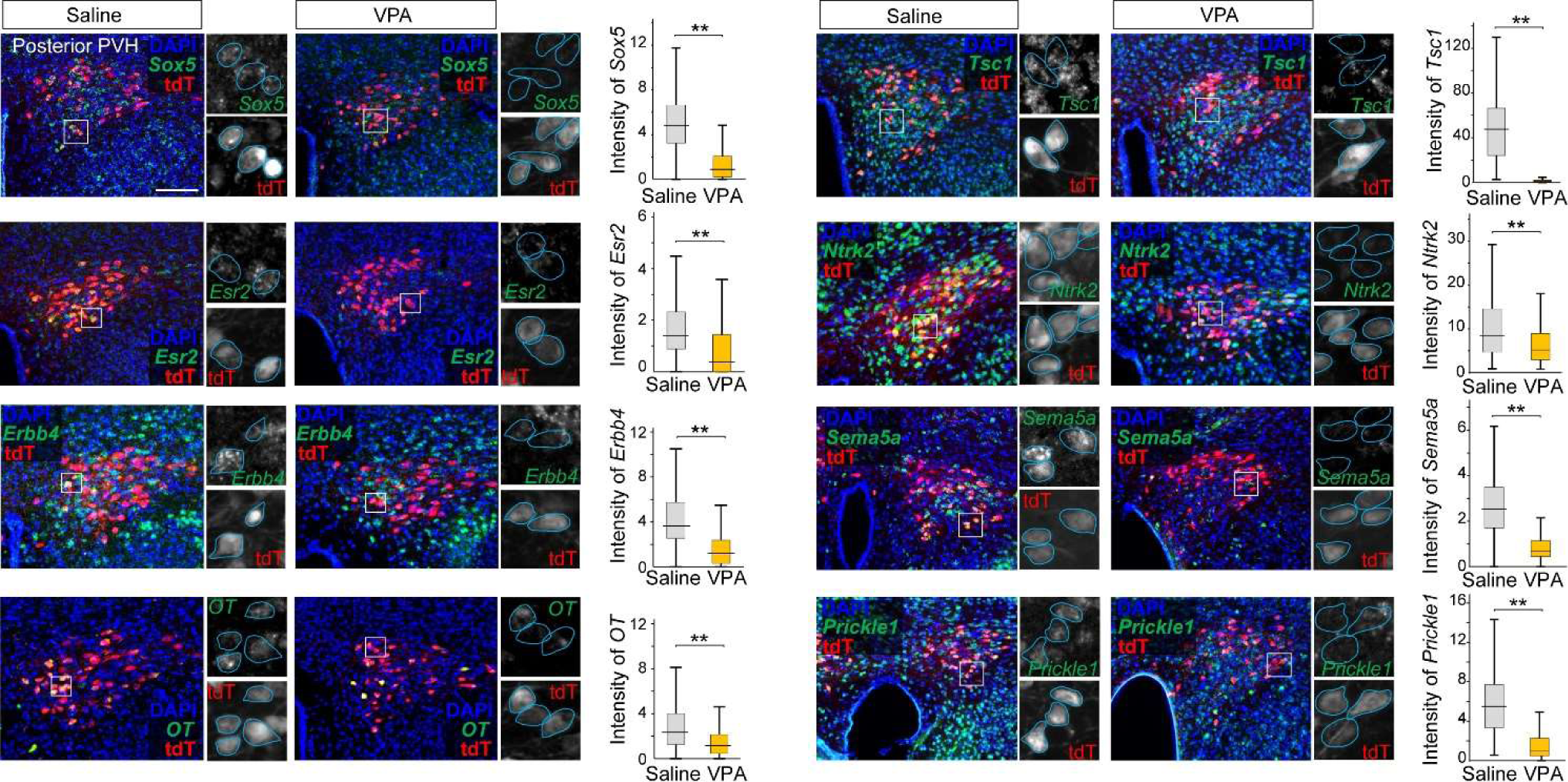
Detection of downregulated DEGs in the parvocellular PVH^OT^ neurons by ISH, related to Fig. 3. Typical coronal sections of the posterior part of the PVH where parvocellular PVH^OT^ neurons exist from *OT-Cre*; *Ai9* mice showing tdTomato and mRNA expressions for indicated genes by ISH. Scale bar, 200 μm. The fluorescent intensity of ISH staining of individual cells for indicated genes is quantified in the graph, in which 100–300 cells from three animals for each condition were analyzed. **, p < 0.01 by the Wilcoxon rank-sum test.

**Extended Data Fig. 6:**
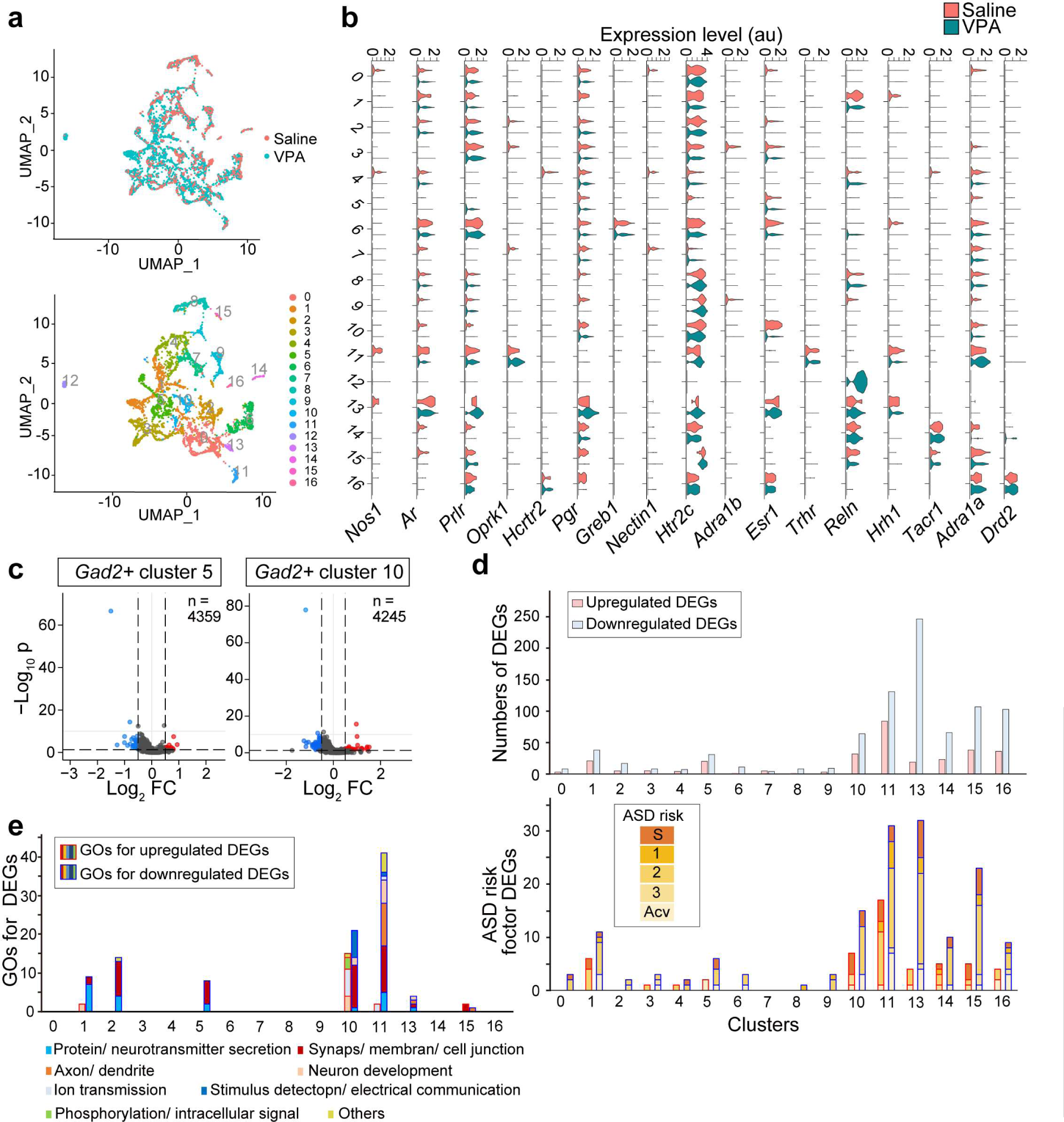
Analysis of DEGs in *GAD2+* neuron clusters, related to Fig. 3. **a**, (Top) UMAP representation of the saline and VPA groups among *GAD2*+ neuron clusters, which are constructed based on Extended Data Fig. 1f. (Bottom) UMAP representations of 17 clusters within the *GAD2*+ neuron clusters. **b**, Violin plots displaying the expression levels of selected marker genes across the 17 clusters identified within the *GAD2*+ neuron cluster. Of note, cluster 12 is specific to the VPA group and characterized by high expression of *Reelin* (*Reln*) gene. **c**, Volcano plots representing the upregulated (red) and downregulated (blue) DEGs in cluster 5 (left) and cluster 10 (right) as in Fig. 2e. **d**, (Top) The numbers of upregulated (red) and downregulated (blue) DEGs within the *GAD2+* clusters. The cluster 12 was excluded from this analysis as this cluster was specific to the VPA group. (Bottom) The number of ASD risk factor DEGs within the upregulated (red frame) and downregulated (blue frame) DEGs. **e**, The numbers of GO terms that show significant association (p < 0.01) with DEGs within the *GAD2+* clusters.

**Extended Data Fig. 7:**
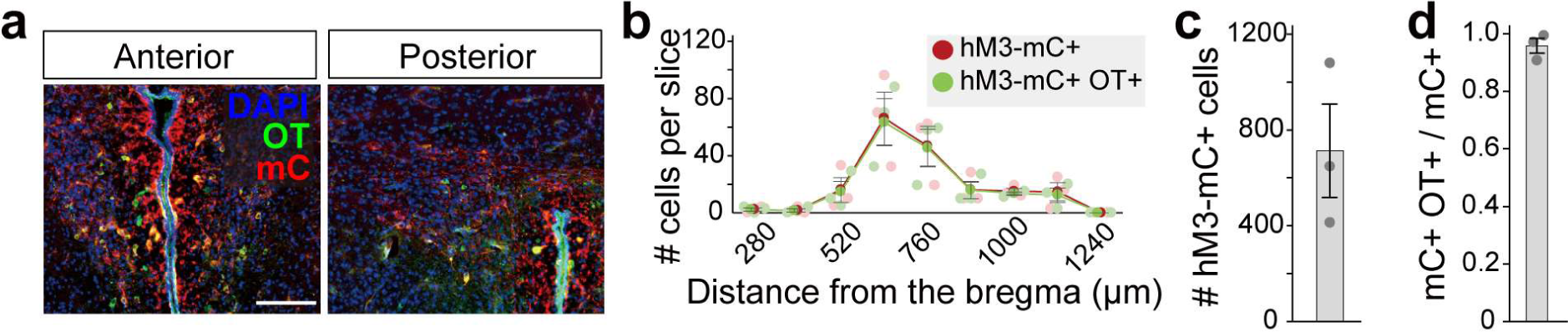
Histochemical analysis of *OT-Cre; stop-hM3* mice, related to Fig. 5. **a**, Typical coronal sections of the anterior and posterior parts of the PVH from typically-developed *OT-Cre*; *stop-hM3* double heterozygous male mice (5 weeks of age) that were prenatally injected with saline. mC, mCherry from the *stop-hM3* allele. Scale bar, 200 μm. **b**, Number of hM3D-mCherry+ cells and hM3D-mCherry+ OT+ dual-positive cells per slice along the anterior–posterior axis of the PVH. The distribution is similar to that observed in *OT-Cre*; *Ai9* mice, suggesting that hM3D is expressed in both magnocellular and parvocellular PVH^OT^ neurons. N = 3. **c**, Number of hM3D-mCherry+ cells, which is comparable to that of the VPA-treated group in Fig. 5i, but smaller than the number of tdT+ cells from the *Ai9* allele in Fig. 1d, presumably because the recombination efficiency of *stop-hM3D* mice is lower than that of *Ai9* mice. **d**, Fraction of hM3D-mCherry+ OT+ cells over hM3D-mCherry+ cells, indicating the high specificity of the transgene expression in PVH^OT^ neurons.

**Extended Data Fig. 8:**
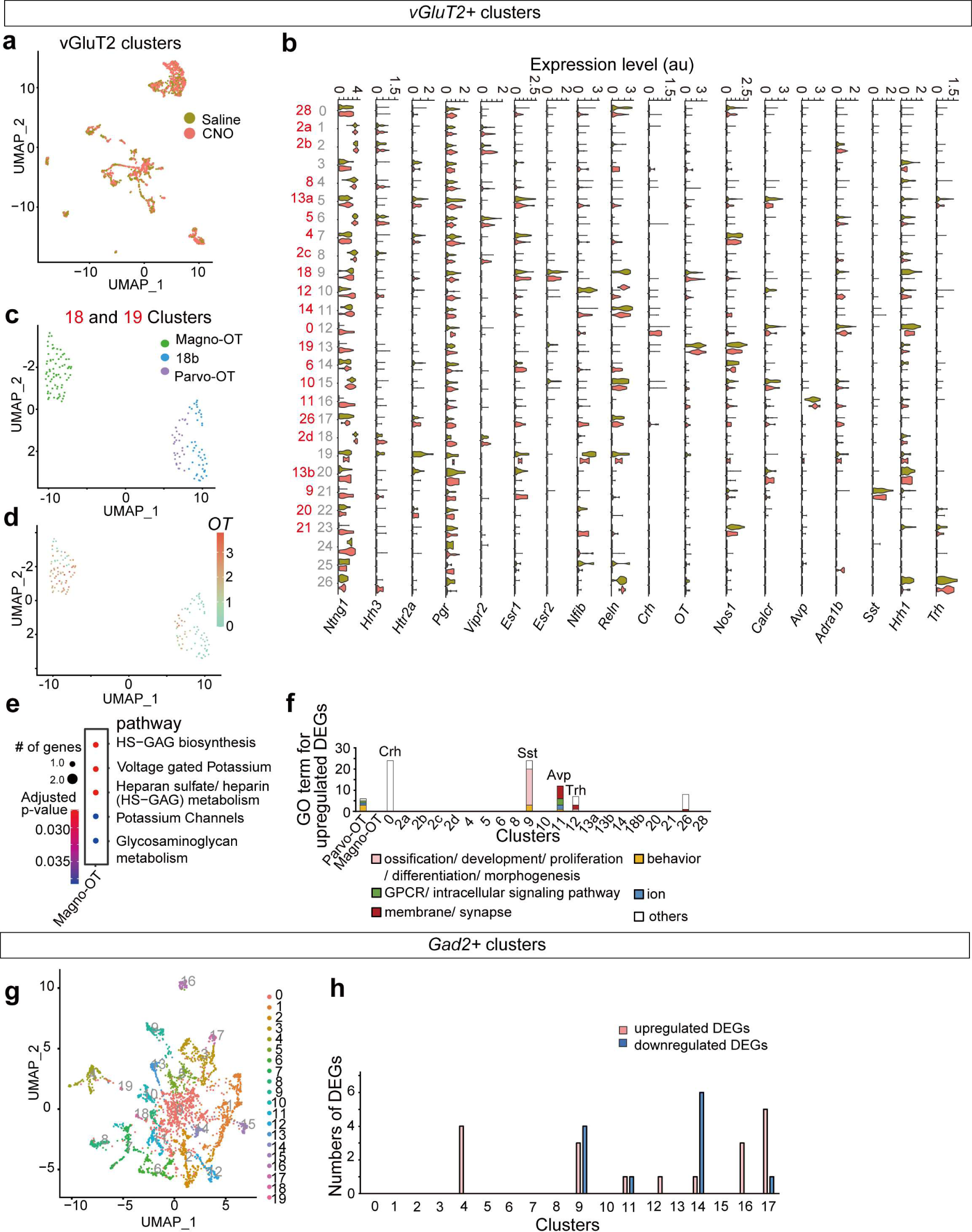
Additional snRNA seq data analyses following neonatal chemogenetic stimulation of OT neurons, related to Fig. 6. **a**, UMAP representation of the saline and CNO groups among *vGluT2*+ clusters (corresponding to Fig. 6b). These two groups were highly intermingled, suggesting that neonatal chemogenetic stimulation did not affect the overall transcriptomic signature. **b**, Violin plots displaying the expression levels of selected marker genes across the 27 clusters identified within the *vGluT2*+ neuron cluster. Based on these marker gene expressions, we manually classified each cluster into one of the *vGluT2*+ clusters in Fig. 2b (shown in red). For 22 out of 27 clusters, we identified the corresponding cluster in Fig. 2b, although the correspondence was not always 1:1. The remaining 5 clusters were not identified in Fig. 2b, presumably due to variance in sampling conditions. **c**, **d**, UMAP showing clusters 9 and 13 (**c**), which correspond to clusters 18 and 19 in Fig. 2b, and *OT* gene expression (**d**), with a color scale showing log-normalized expression. These data indicate that, within this snRNAseq data set, cluster 9 (corresponding to cluster 18 in Fig. 2b) contains two subclusters, one is *OT*+ and the other is *OT*−. We, therefore, classified *OT*+ cluster 9 population as Parvo-OT, while *OT*− cluster 9 population as cluster 18b in the following analysis. **e**, Heatmap representing p-values for pathway analysis using upregulated DEGs in the magnocellular PVH^OT^ neurons. **f**, The number of GO terms that show significant association (p < 0.01) with upregulated DEGs in the *vGluT2*+ clusters (cluster names are based on Fig. 2b). Of note, neonatal stimulation of OT neurons induces non-autonomous effects on *corticotropin-releasing hormone* (*Crh*)+, somatostatin (Sst)+, *Avp*+, *Trh*+ clusters. **g**, UMAP representation of 20 *GAD2*+ neuron clusters. **h**, The numbers of the upregulated (red) and downregulated (blue) DEGs within the top 18 *GAD2+* clusters. Compared with *vGluT2*+ clusters (Fig. 6f), the impact of neonatal OT neuron stimulation was minimal in the *GAD2*+ clusters.

**Extended Data Fig. 9:**
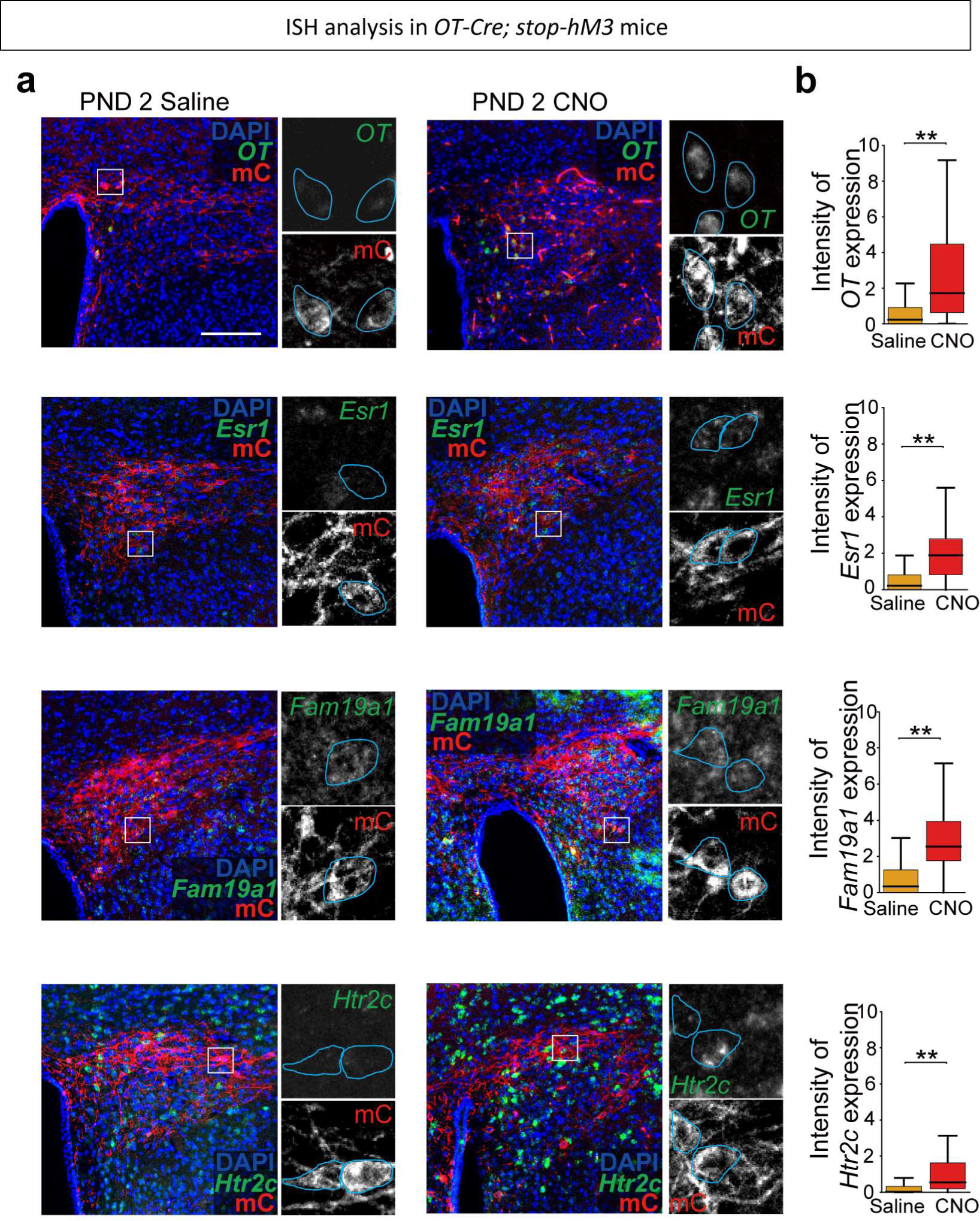
Recovery of mRNA expression in the posterior PVH^OT^ neurons, related to Fig. 6. **a**, Typical coronal sections of the posterior parts of the PVH where parvocellular PVH^OT^ neurons exist from VPA-treated *OT-Cre*; *stop-hM3* male mice (5 weeks of age) that received either saline (left) or CNO (right) at PND 2. These sections were stained with an ISH probe for the *OT, Esr1, Fam19a1,* and *Htr2c* gene (green) together with anti-mCherry (mC) staining (red color, inference of hM3D expression). Sections corresponding to posterior 1000–1120 μm from the bregma were analyzed. Scale bar, 200 μm. **b**, Fluorescent intensity of ISH staining of *OT* mRNA in individual PVH^OT^ neurons. 80–160 cells from three animals for each condition were analyzed. **, p < 0.01 by the Wilcoxon rank-sum test.

## Notes

### Competing Interest Statement

The authors have declared no competing interest.

